# The R2TP chaperone assembles cellular machineries in intestinal CBC stem cells and progenitors

**DOI:** 10.1101/2019.12.19.882712

**Authors:** Chloé Maurizy, Claire Abeza, Valérie Pinet, Marina Ferrand, Conception Paul, Julie Bremond, Francina Langa, François Gerbe, Philippe Jay, Céline Verheggen, Nicola Tinari, Dominique Helmlinger, Rossano Lattanzio, Edouard Bertrand, Michael Hahne, Bérengère Pradet-Balade

## Abstract

The R2TP chaperone cooperates with HSP90 to integrate newly synthesized proteins into multi-subunit complexes, yet its role in tissue homeostasis is unknown. Here, we generated conditional, inducible knock-out mice for *Rpap3* to inactivate this core component of R2TP in the intestinal epithelium. In adult mice, *Rpap3* invalidation caused destruction of the small intestinal epithelium, and death within 10 days. Levels of R2TP substrates were decreased, with strong effects on mTOR, ATM and ATR. *Rpap3*-deficient CBC stem cells and progenitors also failed to import RNA polymerase II in the nucleus. This correlated with p53 activation, cell cycle arrest and apoptosis. Interestingly, post-mitotic, differentiated cells did not display any of those alterations, indicating that R2TP clients are built in actively proliferating cells. Analyses of tissues from colorectal cancer patients revealed that high RPAP3 levels correlate with bad cancer prognosis. Thus, in the intestine, the R2TP chaperone functions in physiologic and pathologic proliferation.

## Introduction

The R2TP complex was first discovered in *Saccharomyces cerevisiae* as an HSP90 co-chaperone (Zhao et al., 2005). HSP90 folds several hundreds of substrate proteins (also called “clients”) into their native, active state (Schopf et al., 2017). The chaperone activity of HSP90 is coupled with its ATPase cycle, and is regulated by a number of co-factors called co-chaperones. These co-chaperones aid client loading and regulate HSP90 ATPase cycle (Schopf et al., 2017). R2TP is an unusual HSP90 co-chaperone for two reasons: first, it is composed of four different subunits; second, it is specialized in quaternary protein folding, i.e. it enables the incorporation of clients into multisubunit complexes (for review (Coulombe et al., 2018)). In mammals, R2TP is composed of a heterodimer between PIH1D1 and RPAP3, which associates with a heterohexamer of RUVBL1 and RUVBL2 (Figue 1A). PIH1D1 is involved in substrate recognition (Hořejší et al., 2014) while RPAP3 recruits the chaperones HSP90 and HSP70 (Pal et al., 2014),(Henri et al., 2018). RUVBL1 and RUVBL2 are related AAA+ ATPases that also have chaperone activity (Zhou et al., 2017),(Zaarur et al., 2015). Multiple contacts between the PIH1D1:RPAP3 heterodimer and RUVBL1/2 heterohexamer allow a regulation of their ATPase activity (Hořejší et al., 2010). Importantly, the RPAP3:PIH1D1 heterodimer is specific to R2TP whereas RUVBL1/2 are also part of other complexes such as the chromatin remodelers INO80 and SRCAP(Jha and Dutta, 2009). In mammals, R2TP associates with a set of six prefoldins and prefoldin-like proteins that are believed to prevent client aggregation. Altogether, R2TP with prefoldins have been termed the PAQosome for Particle for Arrangement of Quaternary structure (Coulombe et al., 2018).

**Figure 1:**
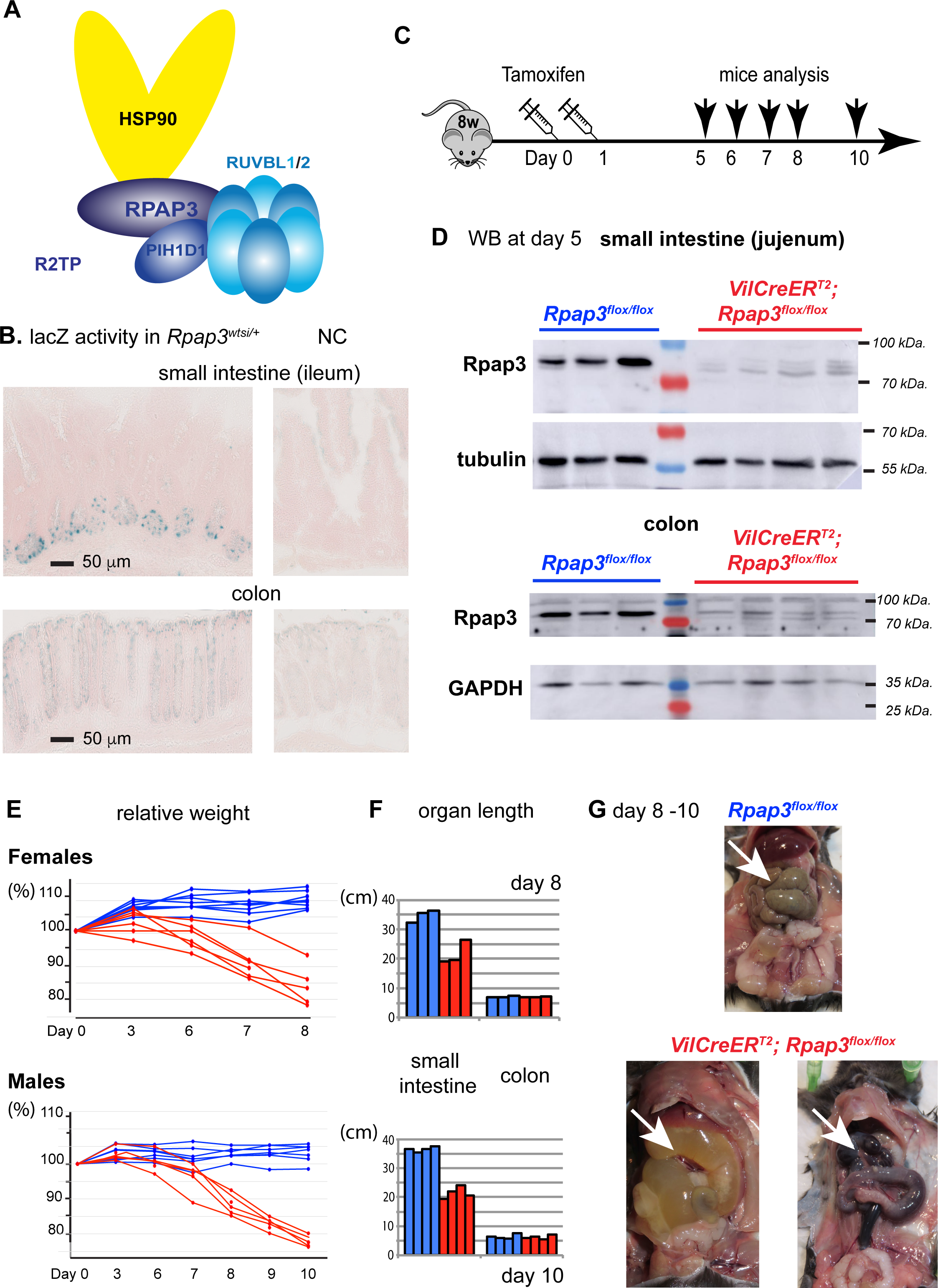
*Rpap3* deletion compromises the small intestine and mouse survival. (A) Schematic representation of R2TP with its four subunits in different tones of blue (RPAP3, PIH1D1 and the RUVBL1/2 hetero-hexamer). RPAP3 is the core subunit that contacts directly HSP90, PIH1D1 and RUVBL1/2. (B) β galactosidase activity in *Rpap3*_*wtsi/+*_ small intestines (top) and colon (bottom), as compared to negative controls (n=2). Scale bars are identical for all images. (C) Schematic representation of the experimental setup. Littermates aged of 8 weeks receive two sequential intra-peritoneal injections of tamoxifen at the indicated time. (D) Depletion of Rpap3 after tamoxifen injection. Western blots were revealed with antibodies against the indicated proteins in extracts of the jejunum and the colonic epitheliums of *Rpap3*_*flox/flox*_ controls (blue) or *VilCreER*_*T2*_; *Rpap3*_*flox/flox*_ animals (red), 5 days after the first tamoxifen injection. Each lane was loaded with the lysate obtained from a single animal. Molecular sizes are indicated on the right. (E) Individual weight variations in females (top panel,) or males (bottom panel) for *VilCreER*_*T2*_; *Rpap3*_*flox/flox*_ animals (red curves, n= 6 females and n=5 males) and *Rpap3*_*flox/flox*_ controls (blue curves, n=8 females and n=7 males). Individual weight was set at 100 for each animal at day 0. (F) Length of small intestines and colons from *VilCreER*_*T2*_; *Rpap3*_*flox/flox*_ mice (red bars) and controls (blue bars). Females were measured at day 8 (top, n=3), and males at day 10 (bottom, n=4). (G) Small intestines (white arrowheads) from *VilCreER*_*T2*_; *Rpap3*_*flox/flox*_ animals were filled with liquid (left panel) or blood (right panel) from day 8 to 10; a control animal is shown above.

The first documented R2TP clients were the small nucleolar ribonucleoparticles (snoRNPs), which are required for the maturation of ribosomal RNAs (Zhao et al., 2008), (Boulon et al., 2008). These particles can be grouped into two families, the C/D and H/ACA snoRNPs, and each family is characterized by a set of four proteins, which assembles with a variety of small nucleolar RNAs. Later on, other R2TP substrates have been identified, including the U4 and U5 spliceosomal snRNPs (Bizarro et al., 2015), (Malinová et al., 2017), (Cloutier et al., 2017), the nuclear RNA polymerases (Boulon et al., 2010), (Cloutier and Coulombe, 2010), and the PI3K-like kinases (PIKKs) mTOR, ATR, ATM, DNA-PK and SMG-1 (Hořejší et al., 2010). R2TP clients thus include multimeric cellular complexes with crucial roles in transcription, ribosome biogenesis and cell growth (Coulombe et al., 2018),(Boulon et al., 2012), yet the role of R2TP in tissue homeostasis has not been studied. Most of our knowledge about R2TP originates from studies in *S.cerevisiae*, mammalian cell lines or *in vitro* studies with recombinant proteins. RPAP3 knock-down does not affect cell line proliferation(Boulon et al., 2010). One study in *D. melanogaster* showed that knock-down of the RPAP3 ortholog Spag hinders early development, but not somatic organogenesis nor homeostasis, suggesting that R2TP could be specialized for stress conditions (Benbahouche et al., 2014). In fact, the role of HSP90 on tissue development is also poorly documented. Constitutive knock-out (KO) murine models showed that one cytosolic paralog of HSP90 was necessary for spermatogenesis while the other one was essential for early development (Grad et al., 2010),(Voss et al., 2000). Conditional models to address the role of HSP90 and R2TP in specific tissues are still missing.

In this study, we chose to address the role of R2TP in the intestine, the most dynamically self-renewing tissue in adult mammals. The intestine has a well-defined architecture which is particularly amenable to study cell proliferation and differentiation (Vermeulen and Snippert, 2014). Briefly, active stem cells called Crypt Base Columnar (CBC) stem cells reside at the bottom of intestinal crypts where they divide to either self-renew, or give rise to progenitors (see schematic Figure 3C). Progenitors migrate through the transient amplifying (TA) compartment to undergo several rounds of division and differentiate. Differentiated cells then migrate towards the tip of the villus, where they ultimately undergo apoptosis and are shed off (Barker, 2013). Progenitors can differentiate into enterocytes, which are responsible for water and nutrient absorption, and compose most of the small intestine epithelium. Alternatively, progenitors differentiate into one of the secretory subtypes. Among these, the goblet cells are the most represented. These cells produce a lubricant protective mucus layer, and their density increases from the duodenum to the colon. In the small intestine, CBC stem cells also generate long-lived Paneth cells (3-6 weeks), which secrete anti-microbial peptides and maintain crypt niche conditions (for review (Barker, 2013)).

To study the role of R2TP in intestinal homeostasis, we generated murine models bearing a conditional knock-out allele of *Rpap3*. Indeed, Rpap3 is central to R2TP, bridging together HSP90, HSP70, PIH1D1 and RUVBL1/2 (Henri et al., 2018),(Martino et al., 2018), (Maurizy et al., 2018),(Muñoz-Hernández et al., 2019). Our work uncovers a crucial role of R2TP in CBC stem cells and progenitors, by coupling the assembly of key cellular machineries with proliferation. In agreement, we found that high RPAP3 expression in the human colorectal cancer (CRC) tissue is associated with a shorter disease-free survival in patients.

## Results

### *Rpap3* is an essential gene in mice

To generate *Rpap3*-deficient mice, we tested different recombined murine Embryonic Stem cells (ES cells) available from the KOMP consortium (for a schematic explanation of the genomic constructs and their products, see Supp. Figure 1A, B). In the *Rpap3*_*wtsi*_ allele, a strong acceptor splice site is introduced before exon 7. The resulting mRNA encodes a Rpap3_wtsi_ protein truncated after the first TPR domain, at amino acid 223, followed by a selection cassette that includes the bacterial β-galactosidase gene (Supp. Figure 1.A, B). Site-specific recombination at the *Rpap3* locus was confirmed by Southern blot in clone H10 (Supp. Figure 1C). Injection of H10 ES cells into blastocysts generated 3 chimeras with germinal transmission. Intercrossing of *Rpap3*_*wtsi/+*_ animals did not yield any homozygous *Rpap3*_*wtsi/wtsi*_ animal, suggesting that this truncation of Rpap3 is homozygous lethal (0/34 pups; Supplemental Figure 1B).

We then took advantage of the β-galactosidase cassette to characterize *Rpap3* promotor activity. In the small intestine of *Rpap3*_*wtsi/+*_ animals, we detected β-galactosidase activity exclusively at the bottom of the crypts, intercalated between *lacZ*_*−*_ cells (Figure 1B). This location could correspond to either CBC stem cells or Paneth cells (see Figure 3C for a schematic representation of small intestinal crypts). In the colon, the signal was weaker and diffuse throughout the epithelium (Figure 1B). These results show that *Rpap3* promoter is active in the intestinal crypts.

### *Rpap3* is required for small intestine maintenance

To generate a conditional knock-out strain, *Rpap3*_*wtsi/+*_ mice were crossed with mice carrying a transgene encoding the Flipase (Flp) (Kranz et al., 2010). In the progeny, excision of the Flp-in cassette produced a *Rpap3* allele with exon 7 flanked by *loxP* sites (Supp. Figure 1A). *Rpap3*_*flox*_ mRNA encodes a protein predicted to be identical to wild-type Rpap3 and indeed, there was no bias in transmission nor any obvious phenotype in *Rpap3*_*flox/flox*_ mice. Excision of exon 7 by Cre recombinase produces a *Rpap3*_*∆7*_ mRNA with a premature termination codon in exon 8. This is far upstream of the wild-type termination codon in exon 17, predicting *Rpap3*_*∆7*_ mRNA degradation by the nonsense-mediated decay pathway (Supp. Figure 1A, B).

To obtain a constitutive deletion of *Rpap3* in the intestinal epithelium, *Rpap3*_*flox/flox*_ mice were crossed with *villin-Cre* (*VilCre*) transgenic mice. *VilCre* transgene encodes a constitutively active Cre that is specifically expressed in all cells of the epithelium of the small and large intestine (El Marjou et al., 2004). We were unable to generate homozygous *VilCre; Rpap3*_*flox/flox*_ animals (0/100 pups; Supplemental Figure 1A), showing that the absence of *Rpap3* in the intestinal epithelium is lethal. To bypass this problem, we turned to an inducible CreER_T2_ under the control of the same promotor *(VilCreER*_*T2*_). This Cre-recombinase is also specifically expressed in the intestinal epithelium, but it needs to be activated by tamoxifen (El Marjou et al., 2004). *VilCreER*_*T2*_; *Rpap3*_*flox/flox*_ animals were healthy and did not present any obvious phenotype. We then injected intraperitonealy two doses of tamoxifen separated by an interval of 24 hours, in eight-week-old mice because the intestine is fully developed at this age (Figure 1C). This yielded an efficient recombination, already detectable one day after the first injection (Supp. Figure 1D). The Rpap3 protein was no longer detected in epithelia from both jejunum and colon, 5 days after the first tamoxifen injection (Figure 1D).

*VilCreER*_*T2*_; *Rpap3*_*flox*/flox_ mice showed no obvious phenotype during the first days following tamoxifen injection but suffered from an important weight loss from day 6 onwards, such that most females and males had to be sacrificed at day 8 or 10 (Figure 1 E). The small intestines from these animals were shorter than those of controls and displayed a massive swelling with a transparent liquid or blood (Figure 1 F, G). Histological analysis revealed no alteration in the small intestine of *VilCreER*_*T2*_; *Rpap3*_*fox/flox*_ mice within the first 6 days (Figure 2A, B red square). At day 7, however, epithelial cells in the villi presented an altered morphology with surface enterocyte disorganization and focal crowding (Figure 2B). At day 8, there was a severe destruction of the small intestinal architecture with villous atrophy and tufts of extruding epithelium (Figure 2B). In contrast, the intestines of the *VilCreER*_*T2*_; *Rpap3*_*flox/+*_ mice did not show any phenotype (Supp. Figure 2 A), suggesting that under normal conditions, one allele of wild-type *Rpap3* is sufficient to maintain tissue homeostasis.

**Figure 2:**
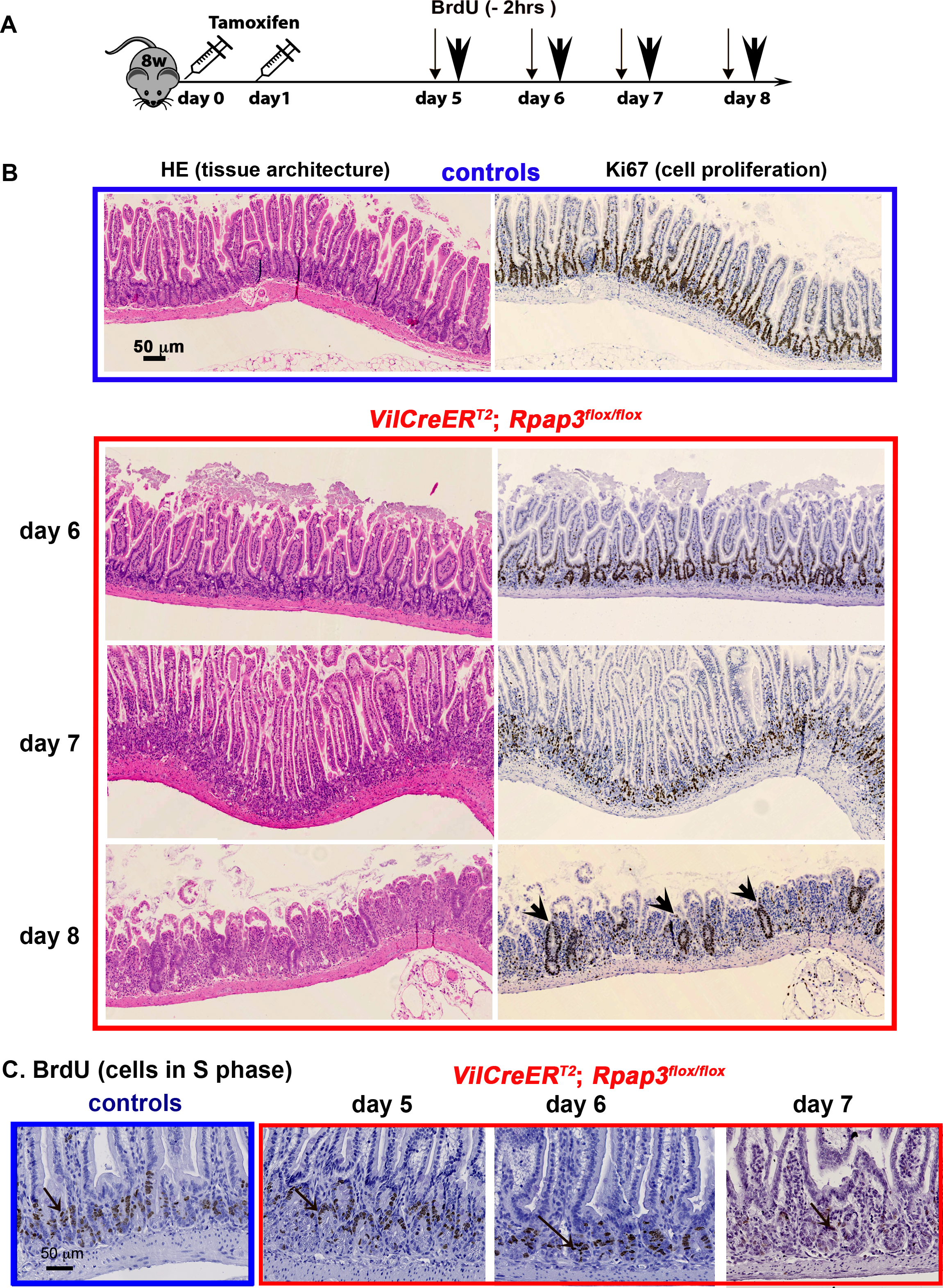
*Rpap3* is necessary for proliferation in the small intestine. (A) Schematic representation of the experimental setting: 8-week-old mice of the indicated genotype received 2 sequential injections of tamoxifen 24h apart, and were analyzed 5 to 8 days after the first injection. 2 hours before each sacrifice, BrdU was injected intra-peritoneally to detect cells in S-phase (thin arrows). (B) Pictures of jejunum tissue sections stained with HE (left) or by IHC with anti-Ki67 antibodies (right panels – Ki67 signal is brown) at different days after the first tamoxifen injection. Black arrowheads indicate remnants of crypt glands with Ki67_+_ cells observed at day 8. Scale bar is shown in control HE panel. Each panel is representative of 8 to 12 animals from three independent experiments. (C) Pictures are jejunum tissue section of controls (boxed in blue) and *VilCreER*_*T2*_; *Rpap3*_*flox/flox*_ animals (boxed in red), stained by IHC with anti-BrdU antibodies (n=3-6 animals/ time point from two independent experiments). Scale bar is shown in control panel.

### R2TP promotes cell proliferation within the epithelium

To understand the basis for this rapid degeneration of the intestinal epithelium, we measured Ki67 expression. Ki67 marks proliferative cells which are, in the intestine, the CBC stem cells and progenitors from the transient amplifying (TA) compartment. Immunostaining revealed a loss of Ki67 in the crypts of *VilCreER*_*T2*_; *Rpap3*_*flox/flox*_ animals starting 7 days after tamoxifen injection (Figure 2B and Supp. Figure 2D for a zoom of the crypt). To directly monitor cell cycling, BrdU was injected into the mouse peritoneum two hours before sacrifice. BrdU is a nucleoside analog incorporated into the DNA during S-phase. BrdU staining was comparable from day 1 to day 6 in the crypts of control and *VilCreER*_*T2*_; *Rpap3*_*flox/flox*_ animals. At day 7 however, there were significantly less BrdU_+_ cells (Figure 2C and Supp. Figure 2E for quantification). This is coherent with the loss of Ki67 staining and confirms cycle arrest in the crypts and the TA compartment. At day 8, the atrophic small bowel mucosa showed remnants of crypt glands with Ki67_+_ cells (Figure 2B, black arrowheads). Such remnants of crypt glands have also been reported upon TCF4 ablation (van Es et al., 2012).

To verify if CBC stem cells were affected, we performed immunostaining for Olfm4, a trans-membrane protein commonly used for CBC identification (van der Flier et al., 2009) (Figure 3A and Supp. Figure 3A). Olfm4 pattern was similar between control and *VilCreER*_*T2*_; *Rpap3*_*flox/flox*_ mice from day 4 to day 6 after tamoxifen injection. At days 7 and 8 however, Olfm4 expression was undetectable in many *VilCreER*_*T2*_; *Rpap3*_*flox/flox*_ crypts (Figure 3A,B and Supp. Figure 3A), in agreement with the loss of Ki67_+_ staining (Supp. Figure 2 C, D). In the crypts, CBC stem cells intercalate between larger differentiated cells, called Paneth cells. These cells are Ki67_-_ and recognizable with the lysozyme marker (Figure 3C, D) or by pink cytoplasmic dots stained by Hematoxylin/Eosin (HE), (Supp. Figure 2 C, D). In marked contrast to CBC stem cells, Paneth cells were detected in a comparable manner in the crypts of control and *VilCreER*_*T2*_; *Rpap3*_*flox/flox*_ mice until day 8 (Supp. Figure 2D and Figure 3D). Altogether, these observations confirmed that loss of *Rpap3* induces the rapid disappearance of proliferating stem cells and progenitors, but not of differentiated Paneth cells.

**Figure 3:**
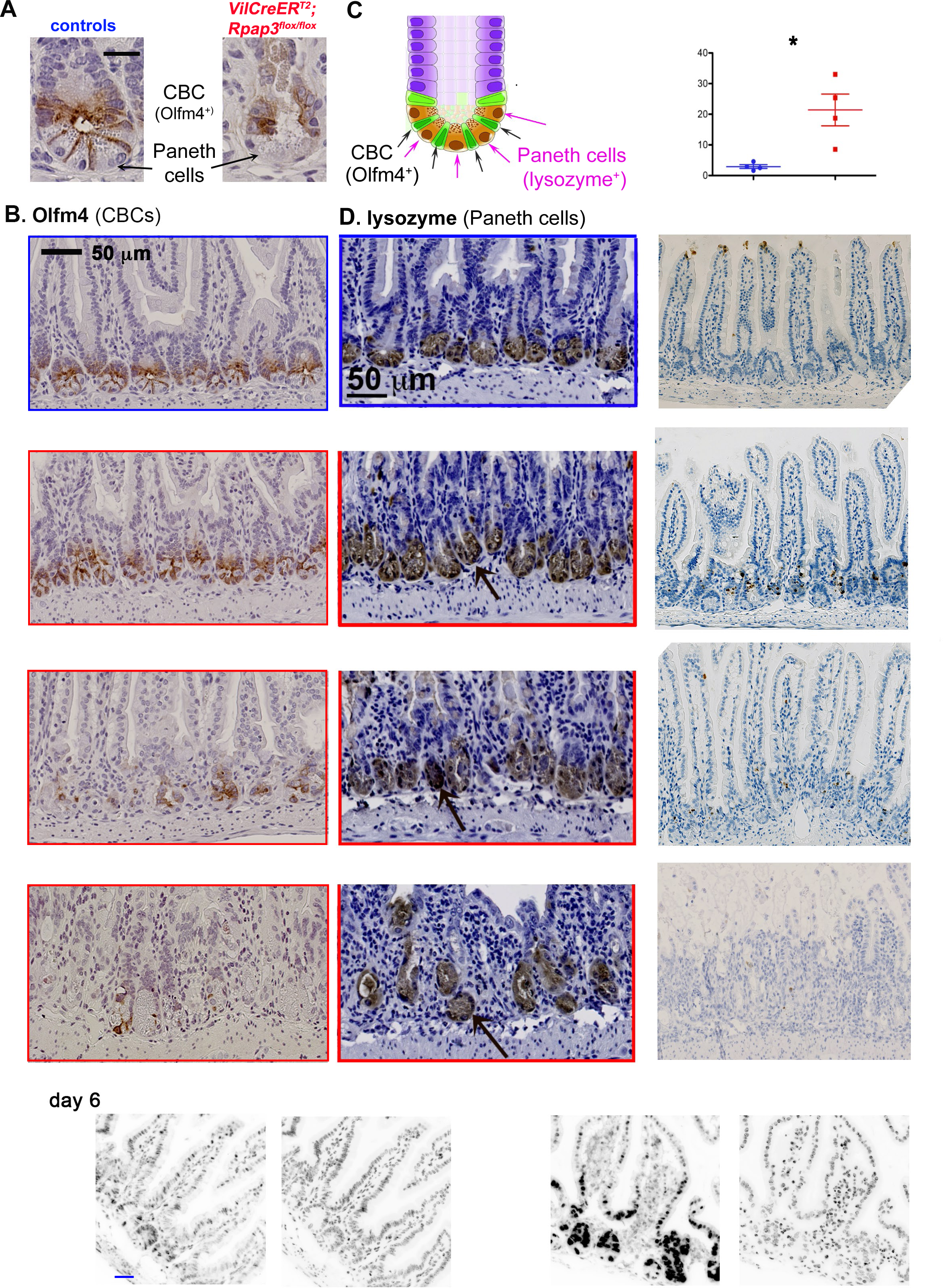
*Rpap3* invalidation induces CBC stem cells loss, p53 stabilization and apoptosis in the proliferative compartment of the small intestine. (A) Micrographs are tissue sections stained for Olfm, in the crypts from controls animals (left), or *VilCreER*_*T2*_; *Rpap3*_*flox/flox*_ animals 7 days after the first tamoxifen injection (right). (B) Staining for Olfm4 in the jejunum from controls (top panel) and *VilCreER*_*T2*_; *Rpap3*_*flox/flox*_ animals 6 to 8 days after the first tamoxifen injection. Scale is identical in all images (n=2 to 4 animals/time point from at least two independent experiments). (C) Schematic representation of a crypt from the small intestine, with CBC stem cells (in green) sandwiched between Paneth cells (in brown) and progenitors forming the TA on top (purple). (D) Micrographs are tissue sections stained for lysozyme, a specific marker of Paneth cells, in the jejunum of controls (top) and *VilCreER*_*T2*_; *Rpap3*_*flox/flox*_ mice from day 6 to day 8 (black arrows). Panels are representative for 2-3 animals/time-point from two independent experiments. The scale bar is identical in all pictures. (E) Micrographs are tissue sections stained for cleaved caspase 3 (cleaved cas3), a marker of apoptosis, in the jejunum. In control animals, cleaved cas3_+_ cells (brown arrows) are mainly detected at the tip of the villi, as a result of epithelium turnover (top panel). In the jejunum from *VilCreER*_*T2*_; *Rpap3*_*flox/flox*_ animals, at day 6 and 7, cleaved caspase 3_+_ cells were detected within the crypts(brown arrows). Panels are representative from 2 to 4 animals/time point, from two independent experiments. The graph on top of the column shows the total number of apoptotic cells at day 6, divided by the surface of the jejunum for each mouse analyzed. Mean values with S.E.M are indicated for each experimental group. Unpaired two-tailed t-test with Welch’s correction indicates significant difference between controls and *VilCreER*_*T2*_; *Rpap3*_*flox/flox*_ animals (p=0, 0384; t=3,538, df=3, n=4). (F) Micrographs are tissue sections stained for p53 by immunofluorescence in controls (left) and *VilCreER*_*T2*_; *Rpap3*_*flox/flox*_ mice (right) at day 6 (n=4). Dapi: nuclei.

Interestingly, we observed cells with round, dense nuclei, resembling apoptotic cells, in the crypts of small intestines deficient for *Rpap3*. We verified this by immunostaining for cleaved caspase 3, a well-established marker of apoptosis. In control mice, few caspase 3 positive cells were visible at the tip of some villi, but we detected significantly more cleaved caspase 3_+_ cells in the crypts of tamoxifen-injected *VilCreER*_*T2*_; *Rpap3*_*flox/flox*_ mice (Figure 3E). These apoptotic cells were observed in the crypts and TA compartment, but not at the tip of villi, as observed in normal epithelia because of cellular turnover. Thus, *Rpap3* deletion eventually induces apoptosis of CBC stem cells and progenitors.

Under stress, cells stabilize p53, which induces cell cycle arrest and/or apoptosis. Since we observed both events, we analyzed its expression. By IF and WB, we saw an increase of p53 at day 6 in the crypts and the TA compartment (Figure 3F and Supp. Figure 3B). Altogether, these results show that inactivation of *Rpap3* affects proliferative cells by inducting p53 expression, cell cycle arrest and apoptosis.

### *Rpap3* stabilizes clients of diverse structural families

Misfolded HSP90 clients are usually degraded _32_, and we thus determined the expression levels of R2TP clients in *VilCreER*_*T2*_; *Rpap3*_*flox/flox*_ animals 6 days after the first tamoxifen injection, when the epithelial architecture was still preserved. R2TP has first been characterized as an assembly factor for the box C/D snoRNPs by recruiting NOP58, one of the four core proteins in these particles (Boulon et al., 2008). Western blot analysis of crypt epithelial cell lysates showed a near two-fold reduction of NOP58 levels in the extracts from *VilCreER*_*T2*_; *Rpap3*_*flox/flox*_ animals, as compared to controls (Figure 4A, quantification in Supp. Figure 4A). PRP8 and EFTUD2 are two core components of the U5 snRNP, a splicing ribonucleoparticle assembled by R2TP in HeLa cells (Malinová et al., 2017),(Cloutier et al., 2017). Both proteins were also reduced by about two-fold (Figure 4A et Supp. Figure 4A, B). HSP90 and R2TP are thought to stabilize clients only before their assembly, and the extreme stability of snoRNPs and snRNPs may thus explain the moderate but consistent decrease observed here for their components (Lemm et al., 2006).

**Figure 4:**
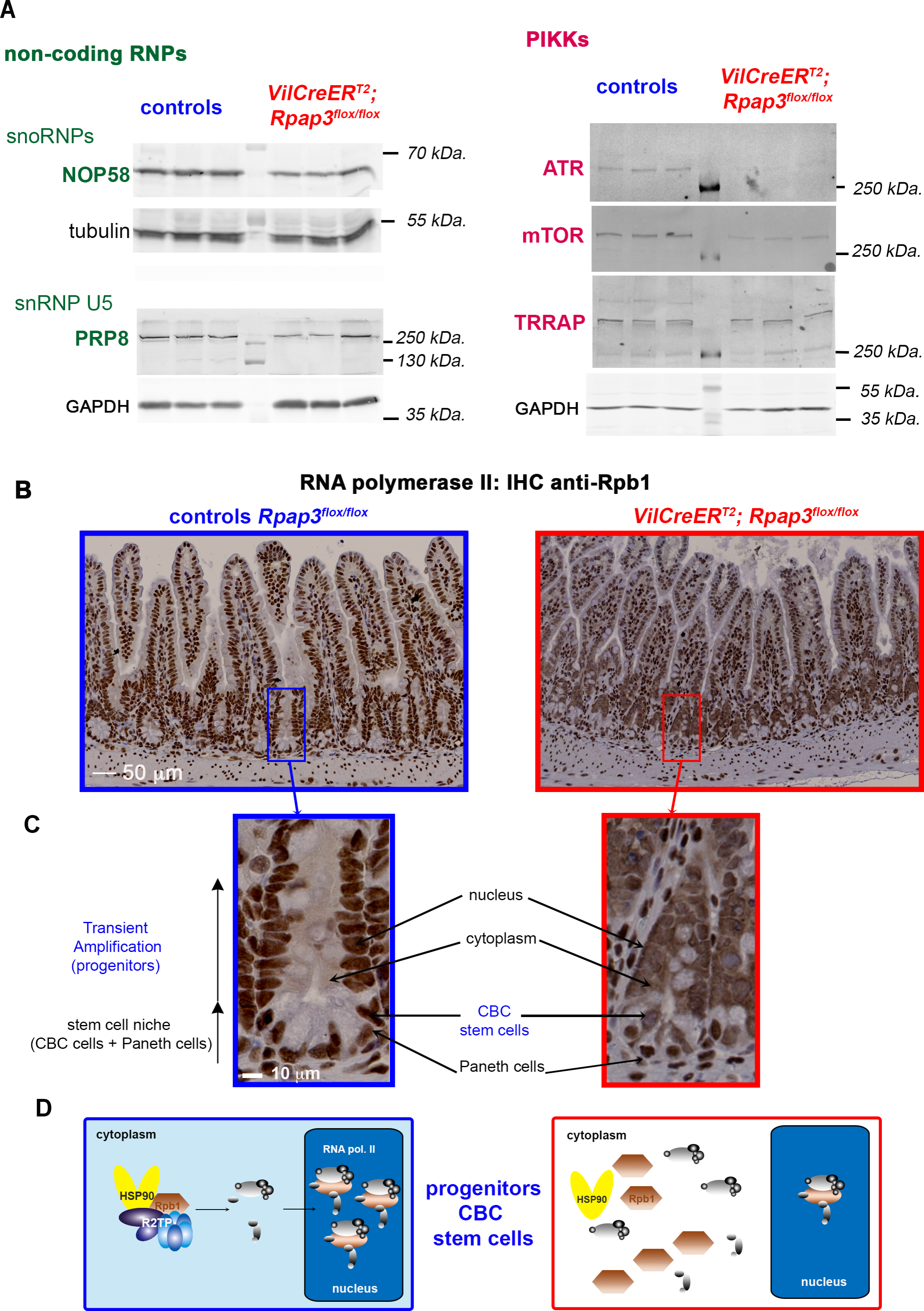
*Rpap3* deletion decreases expression of R2TP clients and leads to cytoplasmic accumulation of RNA polymerase II in intestinal crypts and TA compartment. (A) Western blot analysis of preparations enriched for epithelial crypt cells from the jejunum of animals, sacrificed 6 days after the first tamoxifen injection. NOP58, PRP8, ATR and mTOR and TRRAP were detected with specific antibodies. Tubulin and GAPDH were used as loading controls. Each lane was loaded with the lysate obtained from one animal of the indicated genotype (6 animals per gel). Molecular weights are indicated on the right. See Supplemental Figure 4 for an expanded view of the membranes and signal quantification. (B) Images are tissue sections of small intestines stained by immunohistochemistry (IHC) for Rpb1, the large subunit of RNA polymerase II, from control *Rpap3*_*flox/flox*_ mice (blue frame, left panel), or *VilCreER*_*T2;*_ *Rpap3*_*flox/flox*_ animals at day 6 (red frame, right panel). Note that the staining in stromal cells is nuclear in both wild-type and *VilCreER*_*T2*_; *Rpap3*_*flox/flox*_ animals (stromal cells do not express the Cre), while it becomes cytoplasmic in the mutant epithelium. The scale is identical for both pictures. Panels are representative for n=6 animals from three independent experiments. (C) Magnification of crypts from (B). The scale bar is identical for both images. (D) Schematic interpretation of the IHC in (B, C). In control epithelial cells, R2TP incorporates Rpb1 into RNA PolII, which is then imported into the nucleus. In the absence of Rpap3, neo-synthesized Rpb1 accumulates in the cytoplasm.

In mammals, PIKKs consist of six structurally related proteins (ATR, ATM, DNA-PK, mTOR, TRRAP and SMG1). PIKKs are stabilized by the trimeric co-chaperone TTT (Takai et al., 2007), (Izumi et al., 2012), (Hurov et al., 2010), (Kaizuka et al., 2010) and in murine fibroblasts, TTT recruits R2TP to assemble PIKKs with their partners (Hořejší et al., 2010). We analyzed by Western blot the expression of PIKKs in extracts enriched for crypt cells. We observed a strong diminution of ATR and ATM, the primary sensors of DNA damage, as well as for mTOR, which activates translation and cell proliferation (Figure 4A and Supp. Figure 4 C, D). These results show that R2TP participates in the stabilization of ATR, ATM, and mTOR. In contrast, TRRAP levels did not vary, either because R2TP does not chaperone TRRAP or because it is a very stable protein.

Another R2TP client is Rpb1, the biggest subunit of RNA polymerase II (RNA PolII)(Boulon et al., 2010), (Cloutier and Coulombe, 2010). R2TP incorporates neo-translated Rpb1 within RNA PolII in the cytoplasm, after which it is addressed to the nucleus (Boulon et al., 2010). As expected, IHC for Rpb1 displayed a nuclear staining in control intestines. However, in *Rpap3*-deficient animals, Rpb1 accumulated in the cytoplasm of CBCs and TA progenitors. Interestingly, it remained nuclear in differentiated epithelial cells, including Paneth cells (Figure 4B, C). This revealed that Rpap3 is necessary for the assembly and nuclear import of RNA pol II in the proliferative compartment (Figure 4D).

### *Rpap3* activity correlates with cellular turnover in small intestine and colon

Tamoxifen treated *VilCreER*_*T2*_; *Rpap3*_*flox/flox*_ animals displayed comparable depletion of Rpap3 protein in either small intestines or colons (Figure 1D). Nevertheless, none of the phenotypes affecting the small intestine were observed in the colon: the size of the organs did not vary between control and *Rpap3* KO mice (Figure 1F), nor did their architectural organization, as observed by staining with Periodic Acid Schiff (PAS) or Ki67 (Supp. Figure 5A). To verify whether *Rpap3* deletion affects colonic crypts, we generated *Lgr5-EGFP-IRES-CreER*_*T2*_; *Rpap3*_*flox/flox*_ animals (Figure 5A). In these mice, GFP and the tamoxifen-inducible Cre are both under the control of the Lgr5 promoter, which is only expressed in CBC stem cells (Barker et al., 2007). This driver has a mosaic expression, which allows to analyze Rpap3-deficient and - expressing crypts in parallel. Tamoxifen treated *Lgr5-EGFP-IRES-CreER*_*T2*_*;Rpap3*_*flox/flox*_ mice did not show any visible physiological alteration and could survive for at least 3 weeks.

**Figure 5:**
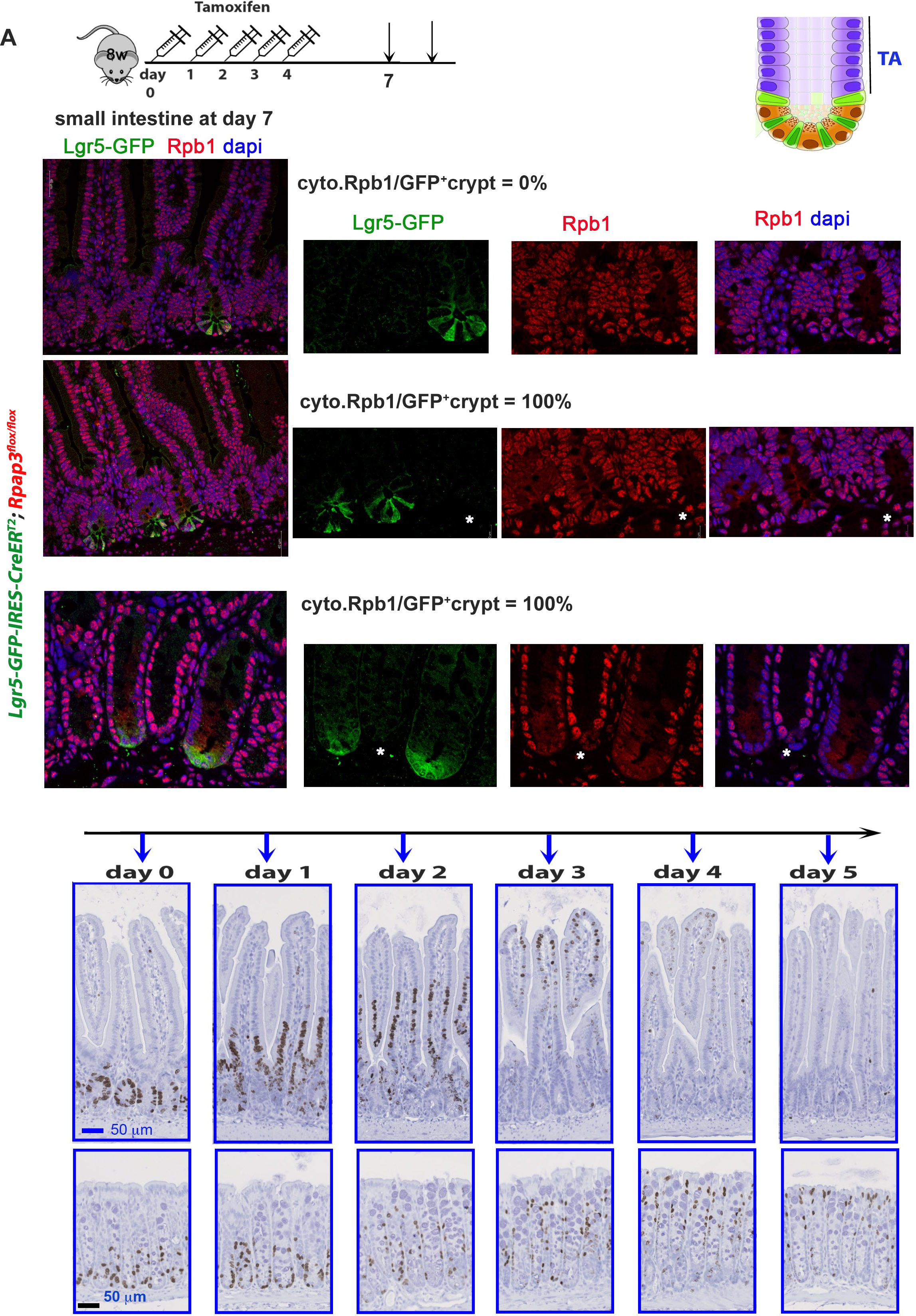
Cytoplasmic accumulation of Rpb1 in *RPAP3* KO crypts correlates with cellular turnover. (A) Schematic representation of the experimental setting. Eight-week-old *Lgr5-GFP-IRES-CreER_T2_; Rpap3_flox/flox_* mice received five sequential intra-peritoneal injections of tamoxifen and were analyzed 7 or 10 days after the first injection. In this genetic model, the Cre is expressed in the Lgr5_+_ CBC stem cells labeled by GFP (see scheme on the right). (B) Images are tissue sections labelled by immunofluorescence with antibodies against GFP (green) and Rpb1 (red), with dapi counter-staining of nuclei (blue). White arrow: GFP+ crypts. Asterisks: GFP-crypts. At day 7, control *Lgr5-GFP-IRES-CreER*_*T2*_; *Rpap3*_*+/+*_ animals do not show detectable Rpb1 signal in the cytoplasm (0/29 GFP+ and 0/96 GFP_-_ crypts; n=2). In the jejunum of *Lgr5-GFP-IRES-CreER*_*T2*_; *Rpap3*_*flox/flox*_ animals (n=3), GFP_+_ crypts show detectable Rpb1 signal in the cytoplasm (50/50). In contrast, adjacent GFP_-_ crypts, which do not express CreER_T2_, do not show cytoplasmic Rpb1 staining (0/136, white asterisk). At day 10, in the colons of *Lgr5-GFP-IRES-CreER*_*T2*_; *RPAP3*_*flox/flox*_ animals (n=2), Rpb1 is detected in the cytoplasm of CBC stem cells and progenitors in the GFP_+_ crypts (white arrow; 11/11). In contrast, Rpb1 was strictly nuclear in the neighboring GFP_-_ crypts (white asterisk, 0/15). Scale bars are identical for matching panels. (C) Images are intestine tissue section of wild-type animals stained for BrdU. Animals received one BrdU injection and were sacrificed at the indicated time point. Scale bars are identical in each row. The experiment was repeated twice (2 to 4 animals/time-point/ experiment).

To analyze the activity of R2TP in the *Lgr5-EGFP-IRES-CreER*_*T2*_*;Rpap3*_*flox/flox*_ mice, we performed a double staining for Rpb1, and GFP to identify crypts with an active Cre. Seven days after the first tamoxifen injection, Rpb1 was localized in the nucleus of GFP_-_ crypts of the small intestine but accumulated in the cytoplasm of CBC stem cells and progenitors from GFP_+_ crypts (100% of GFP_+_ crypts; Figure 5B). In *Lgr5-EGFP-IRES-CreER*_*T2*_; *Rpap3*_*+/+*_ control mice, Rpb1 was nuclear in both GFP_-_ as well as in GFP_+_ crypts (Figure 5B). At day 10, we did not detect any GFP_+_ crypt in the small intestine of *Lgr5-EGFP-IRES-CreER*_*T2*_; *Rpap3*_*flox/flox*_ mice, possibly because recombinant CBC stem cells had been eliminated and replaced by GFP_-_ CBC-derived crypts. In the colon, Rpb1 localization was affected but with a delay, as compared to the small intestine (Supp. Figure 5B). At day 7, Rpb1 was nuclear in GFP_-_ and GFP_+_ colonic crypts from *Lgr5-EGFP-IRES-CreER*_*T2*_; *Rpap3*_*flox/flox*_ mice (Supp. Figure 5C). At day 10, however, Rpb1 accumulated in the cytoplasm of CBC stem cells and progenitors of GFP_+_ crypts (Figure 5B). In colons of control animals, Rpb1 was nuclear in GFP_−_ and GFP_+_ crypts (Supp. Figure 5C). Thus, Rpap3 is also required for Rpb1 assembly in the colon but cytoplasmic accumulation of Rpb1 is detectable with a delay of three days, as compared to the small intestine.

We then decided to compare the cellular turnover in both tissues. For this, we injected the nucleoside analog BrdU as a cell tracer into wild type mice and sacrificed them subsequently at different time points (Figure 5C). Two hours after injection, BrdU was incorporated exclusively by CBC stem cells and progenitors in both the small intestine and the colon (Figure 5C). After one day, BrdU_+_ cells were detectable above the TA compartment in the small intestine, while some remained at the bottom of the crypt. BrdU_+_ cells then migrated to the tip of the villus at day 3, and were eliminated at day 4, in accordance with the kinetics previously described (Tetteh et al., 2016). In contrast, in the colon, migrating BrdU_+_ cells did not reach the tip of the crypt before day 4 and were still detectable at day 5 (Figure 5C). This illustrates a slower cellular turnover in the colon than in the small intestine, correlating with our finding that neo-synthesized Rpb1 accumulates more slowly in the cytoplasm of colonic cells.

### High expression of RPAP3 correlates with poor prognosis in colorectal cancer patients

Our previous findings suggest a strong link between R2TP activity and cell proliferation during intestinal homeostasis. To test if R2TP is also involved in pathogenic proliferation, we took advantage of the CODREAD dataset (available at https://xenabrowser.net/). This showed a significant enrichment of mRNAs encoding RUVBL1, 2 and RPAP3 in human primary colorectal tumors (n= 380) as compared to normal tissue (n=51), in line with a previous report on a smaller cohort (Kim et al., 2013) (Fig. 6A).

**Figure 6:**
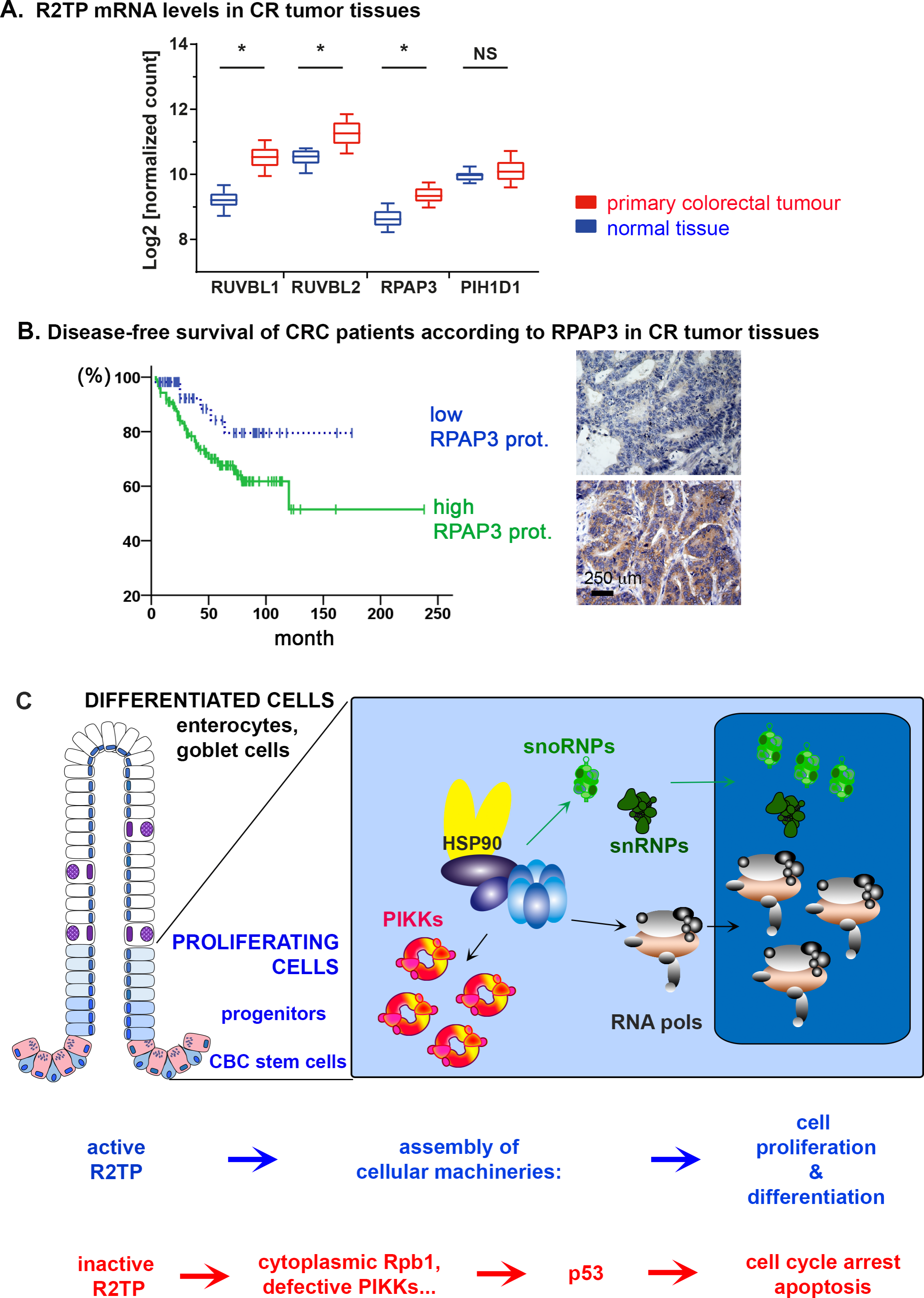
R2TP expression correlates with pathological cell proliferation. (A) The graph depicts transcript levels of R2TP components in human primary colorectal tumor samples (n = 380; COADREAD cohort), as compared to normal solid tissues (n = 51). y-axis: Log2 normalized counts for the indicated transcript. Distributions are presented as box-and-whisker plots (center line: median; box limits, first and third quartiles; whiskers, 10th and 90th percentiles). Statistical significance was determined by one-way ANOVA (asterisk: P < 0.001). (B) Kaplan–Meier analysis of disease-free survival among 177 CRC patients according to the proportion of RPAP3-expressing cells in tumor tissues (C, *P* = 0.037). Solid green line and dashed blue line indicate High and Low proportion of RPAP3-expressing tumoral cells, respectively. Right panels show examples of CRC tissues with low (top) or high (bottom) RPAP3 expression. (C) Proposed model for R2TP activity in the small and large intestine. R2TP assembles cellular machineries such as RNA polymerases, snoRNPs, snRNPs and PIKKs-complexes in CBCs and progenitors in the proliferative compartment (blue cells). Differentiated cells (including Paneth cells from the small intestine crypts, in pink) mostly rely on the complexes assembled during the proliferative phase. A defect in R2TP activity induces client dysfunction, cell cycle arrest and apoptosis *via* p53, and eventually, epithelium degradation.

This prompted us to test the expression of RPAP3 by immunohistochemistry on Tissue Microarrays (TMAs) sections from colorectal cancer (CRC) patients, employing customary anti-RPAP3 antibodies (unfortunately, this antibody detects human RPAP3 but not its murine homolog (Machado-Pinilla et al., 2012)). We analyzed TMAs containing core tissues from patients diagnosed with CRC without pathological evidence of nodal involvement and distant metastasis. One hundred and fifty-seven out of 177 (88.7%) cases expressed RPAP3 in the tumor cell cytoplasm. The proportion of RPAP3-positive cells was in the range of 4-100%, with a mean ± S.E. of 62.6% ± 2.6. To dichotomize the RPAP3 expression level in RPAP3_high_ and RPAP3_low_, an optimal cut-off value of 26% of positive tumor cells was chosen based on the Receiver Operating Characteristic (ROC) analysis (AUC = 0.593). Examples of low and high expression of RPAP3 are shown in Figure 6B. The relationships between RPAP3 expression and clinico-pathological parameters were investigated by Pearson’s χ^2^ test: RPAP3 expression negatively correlated with the tumor stage of CRC (*P* = 0.049) (Supplemental Table 1, 2).

Thirty-nine out of 123 (31.7%) patients with RPAP3_high_ tumors and 6 out of 54 (11.1%) patients with RPAP3_low_ tumors had disease relapse. Analysis of Kaplan–Meier curves showed that patients with RPAP3_high_ tumors had a lower Disease-Free Survival (DFS) rate than patients with RPAP3_low_ tumors (*P* = 0.037; Figure 6B). Multivariate analyses of DFS adjusted for other prognostic factors confirmed that RPAP3 expression was a significant prognostic parameter influencing disease relapse (HR = 2.7: 95% CI, 1.2-6.5; *P* = 0.023) (Supp. Table 3), but not the overall survival of patients. These results provide evidence that high RPAP3-expression levels in CRC tissues are associated with poor patient prognosis.

## Discussion

### R2TP inactivation leads to client dysfunction, p53 activation and apoptosis

Since its discovery in 2005, structural studies of the R2TP chaperone have provided substantial knowledge (Zhao et al., 2005), (Coulombe et al., 2018). It is now clear that RPAP3 is a central factor within R2TP. It recruits the chaperones HSP70/90 *via* its TPR domains, PIH1D1 through a short peptide domain and the AAA+ ATPases RUVBL1/2 *via* its C-terminal domain (Henri et al., 2018), (Martino et al., 2018), (Maurizy et al., 2018), (Muñoz-Hernández et al., 2019). Nevertheless, how R2TP ultimately assembles neo-synthesized proteins into active complexes is still unknown. Most of the R2TP clients have been identified in mammalian cell lines (Cloutier and Coulombe, 2010), (Boulon et al., 2008), (Boulon et al., 2010), (Hořejší et al., 2014), (Cloutier et al., 2017), (Malinová et al., 2017). For several of them, we validate here their dependence on RPAP3 in a mammalian tissue. This includes PRP8, part of the splicing snRNP U5, and NOP58, a core component of box C/D snoRNPs. Another class of R2TP substrates are PIKKs (Hořejší et al., 2010). Several studies have documented that these large proteins are stabilized by a trimeric co-chaperone called TTT, which itself interacts with R2TP (Hurov et al., 2010), (Kaizuka et al., 2010), (Izumi et al., 2012), (Takai et al., 2010). We show here that mTOR, ATR and ATM depend on *Rpap3* in intestinal crypts. Finally, a major R2TP client is RNA polII. This key enzyme is assembled in the cytoplasm and its Rpb1 subunit remains cytoplasmic when assembly is deficient (Boulon et al., 2010). The observed cytoplasmic accumulation of Rpb1 in *Rpap3* knock-out cells thus likely results from a defective assembly of RNA polII.

Invalidation of *Rpap3* triggers p53 stabilization and cell cycle arrest, ultimately leading to apoptosis and destruction of the intestinal epithelium (Figure 6C). Transcription inhibition is well known to activate p53 (Ljungman, 2000). In addition, while neither mTOR nor ATM loss affect intestinal homeostasis (Sampson et al., 2016), (Brandt et al., 2018), (Barlow et al., 1996), defective ribosome biogenesis or ATR inactivation both induce a degradation of the intestinal epithelium through p53 activation, similar to the one described here (Stedman et al., 2015), (Ruzankina et al., 2009). The alterations that we observed for snoRNPs, ATR and RNA PolII might thus concur to induce the phenotype observed in the *Rpap3* knock-out intestine.

### R2TP is active in the proliferative compartment

We found that differentiated cells in the small intestine are barely affected by *Rpap3* removal, in contrast to the rapidly dividing CBC stem cells and TA progenitors. Indeed, Rpb1 accumulates in the cytoplasm of proliferating cells, while it remains fully nuclear in the differentiated cells of the epithelium. This correlation between proliferation and sensitivity to *Rpap3* removal further extends to the colon. Rpb1 also accumulated in the cytoplasm of colonic recombinant crypts, but this occurred several days later than in the small intestine, in line with its slower proliferation rate. Highly proliferative cells appear to be very sensitive to Rpap3 depletion. We thus propose that basic cellular machineries are assembled by R2TP in the proliferative compartment: CBC stem cells and progenitors.

Several additional data support this model. First, β-galactosidase knock-in models showed that *Rpap3* mRNA is expressed in the crypts where CBCs reside. Second, some R2TP clients are themselves preferentially active in the intestinal proliferative compartment: (i) under homeostatic conditions, ATR is required to signal defective DNA replication in the intestinal proliferative compartment (Ruzankina et al., 2007); (ii) markers of RNAPolI activity have been detected preferentially in the crypts and TA compartment with decreasing levels along the crypt-villus axis (Stedman et al., 2015); (iii), a recent transcriptomic analysis along the crypt-villus axis detected mRNAs encoding ribosomal, splicing and transcription components at the very bottom of the villus, but not along the villus or at the tip (Moor et al., 2018). All this supports a model in which ribosomal biogenesis, transcription machineries and PIKK complexes are built in the rapidly proliferating stem cells and progenitors. These assembled complexes would suffice for the relative short lifespan of differentiated cells, until they are shed off the villus (Figure 6C).

In Drosophila, the RPAP3 ortholog, Spag, is required for ovarian Germline Stem Cell (GSC) maintenance (Chen et al., 2017). Ovarian germline stem cells divide asymmetrically to self-regenerate and give rise to an oocyte. Knock-down of *Spag* induces a loss of GSC but not of the non-dividing somatic cells in the ovary. This exemplifies another case where RPAP3/Spag affects dividing stem cells, and suggests a general role of R2TP in proliferation, especially of active stem cells, across Metazoans.

### R2TP is linked with pathological cell proliferation

HSP90 can sustain tumor growth by folding numerous proteins, some of which are involved in oncogenic signaling pathways. In addition, an enhanced stability of the HSP90 interactome has been observed in some tumors and called the “epichaperome” (Rodina et al., 2016). All this explains why some tumor cells are particularly sensitive towards HSP90 inhibition (Fierro-Monti et al., 2013), (Echeverría et al., 2019), (Pillarsetty et al., 2019). In a mouse model, invalidation of a cytoplasmic HSP90 paralogue (only essential for spermatogenesis (Grad et al., 2010)) diminishes lung metastasis (Vartholomaiou et al., 2017). Yet, in clinical trials, HSP90 inhibitors have been deceptive, showing strong cytotoxicity. This has been attributed to the broad range of HSP90 substrates, which also includes tumor-suppressors (Vartholomaiou et al., 2016), (Suman Chatterjee and Timothy Burns, 2017). To bypass this problem *Neckers et al.* suggested to target HSP90 co-chaperones that are specific for a given class of clients (Neckers and Workman, 2012). We propose to consider R2TP for alternative strategies, as a role for R2TP in intestinal carcinogenesis is supported by several lines of evidences: (i) a higher expression of mRNAs encoding R2TP subunits in colorectal tumors, as compared to matching controls; (ii) high RPAP3 protein levels in biopsies of CRC patients with poor diagnostic, (iii) a strong dependency on RPAP3 in the proliferative compartment, (iv) the induction of p53 upon R2TP inactivation. R2TP is interesting not only as a therapeutic target but also for diagnostic purpose, since R2TP components RPAP3 and PIH1D1 were both identified as part of the epichaperome of HSP90-sensitive tumours (Rodina et al., 2016).

In conclusion, we show that the R2TP chaperone plays a crucial role in the homeostasis of the intestinal epithelium, by regulating proliferating stem cells and progenitors.

## Data and sample availability statement

Mouse sperm for the different strains is available upon request to. Transcriptomic analysis of R2TP subunits in human samples (Figure 6A) used the UCSC Xena platform for public and private cancer genomics data visualization and interpretation https://xenabrowser.net/ (Goldman et al., 2018).

The data concerning the patient biopsies (Figure 6B) are not publicly available because the database contains sensitive information that could compromise patient privacy.

Details about the experimental procedures are available upon request to.

## Acknowledgments

We thank Solange Morera and EMBL facility for help with anti-RPAP3 antibody generation. The ES cells used for this research project were generated by the trans-NIH Knock-Out Mouse Project (KOMP) and obtained from the KOMP Repository (www.komp.org). NIH grants to Velocigene at Regeneron Inc (U01HG004085) and the CSD Consortium (U01HG004080) funded the generation of gene-targeted ES cells for 8500 genes in the KOMP Program and archived and distributed by the KOMP Repository at UC Davis and CHORI (U42RR024244). We thank the CIGM team in Institut Pasteur for technical support in microinjection experiments and animal husbandry. We thank the in-house animal facility, RAM, MRI for their excellent support. We acknowledge the “Réseau d’Histologie Expérimentale de Montpellier” (RHEM) facility supported by SIRIC Montpellier Cancer Grant INCa_Inserm_DGOS_12553, the European Regional Development foundation and the Occitanian Region for processing our animal tissues. We thank the SIRIC Montpellier Cancer Grant INCa_Inserm_DGOS_12553. The work was supported by La Ligue Nationale Contre le Cancer (équipes labelisées to EB, PJ) and the INCa grants PLBIO 2016-161 to DH, EB, MH, BPB and PLBIO 2018-158 to PJ. CM and CA Ph.D. fellowships were from la Ligue Nationale Contre le Cancer.

## Authors contributions

C.M., C.A., V.P., M.F., C.P., J.B., F.G., C. V., B.P.B. performed experiments, F.L. generated murine model, N.T. and R.L. provided human TMA and analysis, D.H. analysed data, F.G., P.J. and D.H. advised on the work and commented critically on the manuscript, C.M., E.B., M.H., B.P.B. conceived the study, designed the experiments, analysed the data and wrote the manuscript. B.P.B. supervised the research. All authors approved the content of the manuscript.

## Methods

### Human colorectal sample collection and analysis, and Tissue Microarray (TMA) construction

A total of 177 CRCs were collected from patients surgically treated at the University “G. D’Annunzio”, Chieti, Italy, between 1996 and 2010. Only the CRC patients who did not receive adjuvant systemic therapy were included in the study. The median follow-up was 53 months (range 3-238 months). During the follow-up, 25.4% of CRC patients (45 out of 177) had a disease relapse, while deaths were observed in 18.6% of CRC patients (33 out of 177). Tumor stage was determined according to the American Joint Committee on Cancer (AJCC) TNM staging system (8_th_ edition). Histologically, each CRC case was graded according to the criteria of the WHO classification of tumors of the digestive system (4_th_ edition). Patients and tumor characteristics are summarized in Supplemental Table 1&2. The study was reviewed and approved by Institutional Research Ethics Committee and written informed consent was obtained from all patients.

TMAs were constructed by extracting 2-mm diameter cores of histologically confirmed neoplastic areas from the 177 CRC cases using a manual Tissue Arrayer (MTA, Beecher Instuments, WI), as previously detailed (Lattanzio et al., 2012).

### Mice generation and treatments

Mouse experiments were performed in strict accordance with the guidelines of the European Community (86/609/EEC) and the French National Committee (87/848) for care and use of laboratory animals and comply the ARRIVE guideline. *Rpap3*_*wtsi/+*_mice were generated from ES cells generated by the trans-NIH Knock-Out Mouse Project (KOMP) and obtained from the KOMP Repository (www.komp.org). Genotyping was performed by PCR amplification using primers F5, R6, on the mouse tail genomic DNA (gDNA – Supplemental Figure 1A). Mice were bred in an SOPF animal facility and maintained during the experiment in an SPF animal facility. Naive mice were minimum 6 weeks old and are euthanized by CO2 and isoflurane. To activate the CreER_T2_, controls and animals of interest received two intra-peritoneal (IP) injection of 2mg tamoxifen each. For BrdU incorporation assays, mice were intraperitoneally injected with 100 µg Bromodeoxyuridine (BrdU) per gram body weight. The entire small intestine (cut in 3 parts) and colon were flushed with PBS, then with neutral buffered formalin (4% formaldehyde) and fix in 24h, dehydrated, and embedded in paraffin.

### Histology and immunostainings

*For histological analysis*, tissue sections (4μm thick) were deparaffinized and rehydrated. They were stained with hematoxylin and eosin (HE), Periodic Acid Shiff staining (PAS) for preliminary analysis.

***For immunostainings***, tissue sections were deparaffinized, rehydrated and subsequently subjected to heat induced antigen retrieval by immersing them, depending on the antibody, either in a water bath with a sodium citrate buffer (pH 6) or an EDTA buffer (pH 9). Immunohistochemistry was performed using a Dako autostainer (Dako, Glostrup, Denmark). After neutralization of the endogenous peroxidase activity, the sections were incubated with the primary antibodies. Antibody was visualized using the Envision® system (Dako). Diaminobenzidine (Dako) was used as the chromogen and the sections were lightly counter-stained with hematoxylin. Slides were prepared by RHEM platform and visualized with a NanoZoomerslide scanner controlled by the NDP.view software (MRI platform).

***For p53 immunofluorescence***, paraffin-embedded tissue was cut into 3-µm-thick sections, mounted on slides, then dried at 37°C overnight. It was performed on the Discovery Ultra Automated IHC staining system from Roche Ventana. Following deparaffination with Discovery EZ Prep solution at 75°C for 24 minutes, antigen retrieval was performed at 95°C for 16 minutes with Discovery CC1 buffer. After blocking with (TBS-10%goat serum-5%BSA-5%milk-0, 3%triton), the slides were incubated after rinsing at 37°C for 60 minutes with a rabbit anti-P53 antibody (Leica, P53-CM5P-L, 1∶250). Signal enhancement was performed using the OmniMap anti-Rabbit HRP (Roche, 760-4457) then Cy5 Kit (Roche, 760-238). Slides were then counterstained for 8 minutes with DAPI and manually rehydrated before coverslips were added.

***For Rpb1/GFP immunofluorescence***, antigen retrieval with 1mM EDTA was performed for 30 min at 99°C. Slides were blocked with (5%goat serum-PBS-0,3%triton X-100) for 1h at RT then with primary antibodies overnight at 4°C. Samples were incubated with Dapi+ Secondary antibodies solution for 1h at RT. Slides were mounted with coverslip and ProLong Gold Antifade Mountant.

***For immunohistochemistry (Rpb1, Lysozyme, Olfm4***), tissue slides were deparaffinized and rehydrated by performing the following washes: xylene, ethanol and dH_2_O. Tissue slides were incubated with 10 mM sodium citrate pH6.0 (T0050, DiaPATH) for 20 min at 100°C or Tris10 mM-EDTA 1mM pH9.0, for antigen retrieval, depending on the antibody. Endogenous peroxidases were inactivated with PBS-0,3% hydrogen (#H1009, Sigma). After blocking in 2.5 % blocking serum-5% BSA-5% non-fat milk for 30 min at room temperature, tissue slides were incubated with primary antibody overnight at 4°C. Then corresponding secondary antibody reagents (ImmPRESS_TM_ kit, Vector Laborarories), directed against rabbit, rat or mouse antibodies were used for detection.

Incorporation of BrdU in proliferating intestinal epithelial cells was detected using an anti-BrdU antibody (Biolegend, 1:100) after deparaffinization of the tissues, antigen retrieval in citrate buffer, as described above, and DNA denaturation using 2NHCl for 1H at 37°C followed by an incubation in 0.1M borax buffer pH9. Revelation was performed using the Avidin/Biotin Vectastain System kit (Vectorlab, USA) according to the protocol.

All antibodies used for immunostainings are described in Supplemental Table 4 and validated by the manufacturers, except for 19B11 (Boulon et al., 2010), (Machado-Pinilla et al., 2012).

#### β-galactosidase activity

Small intestines and colons cryo-preserved were cut to slices of 10 μm thickness. Samples were treated with 0.5% glutaraldehyde for 10 min at room temperature, followed by 24h incubation at 37°C in a staining solution containing 1 mg/ml X-gal, 5 mM potassium ferricyanide (K3Fe(CN)6), 5 mM potassium ferrocyanide (K4Fe(CN)6) and 2 mM MgCl2, 0.1% Triton in PBS.. Samples were washed in PBS for 5 min, counterstained with Nuclear Fast Red and briefly rinsed with dH2O before mounting on aqueous mounting agent Aquatex (108562; Merck).

### Microscopy and imaging

Histological slides were scanned using Nanozoomer 2.0 HT scanner with a 40x objective, and visualized with NDP.view2 viewing software (Hamamatsu). Fluorescent images were acquired on a brightfield microscope (Leica) using Metamorph software or inverted Confocal SP5 (Leica) using the Leica LAS AF software. Images were processed with ImageJ. Images were assembled and adjusted with Adobe Photoshop/ Illustrator.

### Isolation of epithelial cells from the intestine

Small intestines and colons were isolated and flushed with cold PBS. Colon and the 3 intestine fragments were cut open length-wise, then kept in 10 ml cold wash buffer (PBS, 2% FBS and antibiotics). Tubes were shaken several times by hand to wash the fragments and put into 15 ml falcon tube containing 10 ml CE buffer (PBS, 1% BSA, 1mM DTT, 1mM EDTA, 5, 6 mM glucose). Tubes were placed on a vertical shaker at 37°C for 30 min. After removing tissue, cells were collected by centrifugation at 1410 rpm for 7 min. They were washed in 10ml wash buffer. Collected cells were split in different tubes and stored at −80°C.

To enrich for crypt-cells portion, before the incubation with the lysis buffer, the samples are scrapped with a slide to remove the villi part. These villi cells are collected by pipeting in 1 ml of cold PBS. The rest of organ is incubated with 10 ml lysis buffer to dissociate crypt cells.

### Western blots

Proteins were prepared from cell preparation by lysis in cold RIPA buffer (50 mM Tris pH 8, 150 mM NaCI, 1% NP40, 0.5% deoxycholate) supplemented with inhibitors. Protein lysates were separated by gradient 4%-15% SDS-PAGE (BioRad) and transferred to a nitro-cellulose membrane (Amersham Protran 0.2µm NC). Membranes were blocked with 2, 5% non-fat milk (w/v) in TBS with 0.05% Tween and incubated with the appropriate primary antibodies at the appropriate dilution and followed by incubation with secondary antibodies conjugated to a fluorophore. Fluorescence was revealed using a scanner (Amersham Typhoon).

Primary antibodies from Cell Signaling: ATM D2E2 (CST 2873S) rabbit at 1/1000, ATR E1S3S (CST 13934S) rabbit at 1/1000, mTOR (CST 2972S) rabbit 1/1000, p53 1C12 (CST2524S) mouse 1/1000; PRP8 (sc-30207, Santa Cruz) rabbit 1/200; from ABCAM: EFTUD2 (ab72456) rabbit 1/2000, GAPDH (ab8245 6C5) mouse 1/10000; and from Sigma: NOP58 (HPA018472) rabbit 1/100, RPAP3 (SAB1411438) rabbit 1/1000, Tubulin (I2G10) mouse 1/500, TRRAP 2TRR-1B3 (MABE1008) mouse 1/1000. All antibodies have been validated by manufacturers.

### DNA extraction

Samples are incubated in 100 µl of alkaline lysis buffer (NaOH 25mM, EDTA 0,2mM) with at 92°C for 20 min. Then, they are mixed with 100 µl of neutralizing reageant (Tris-HCl-pH 5, 40mM). 2 µl were used for PCR. PCR products are respectively 400bp and 300bp for floxed and wild type allele.

### Southern blot

Briefly, ES cells were incubated o/night at 65°C in 100mM Tris HCL pH8.8 - 5 mM EDTA - 0.2% SDS - 200 mM NaCl with 0, 1 mg/ml proteinase K in a humid chamber. On the following morning, the lysates were extracted twice with phenol pH= 7, 0-8, 0 and once with chloroform and precipitated with NaCl/EtOH. Following recovery, 10 – 15 μg genomic DNA were digested for 24h to 48h with appropriate highly concentrated restriction enzymes (40-50 U/μl – 10 U/μg gDNA). The analysis was performed by Southern blot as described in (Brown, 2001).

### PCR for genotyping

Initatial denaturation:95°C (2min), Denaturation 95°C (30sec), Annealing 55°C (45sec), Extension 72°C(45sec), Final extension: 72°C (2min) 35 cycles.

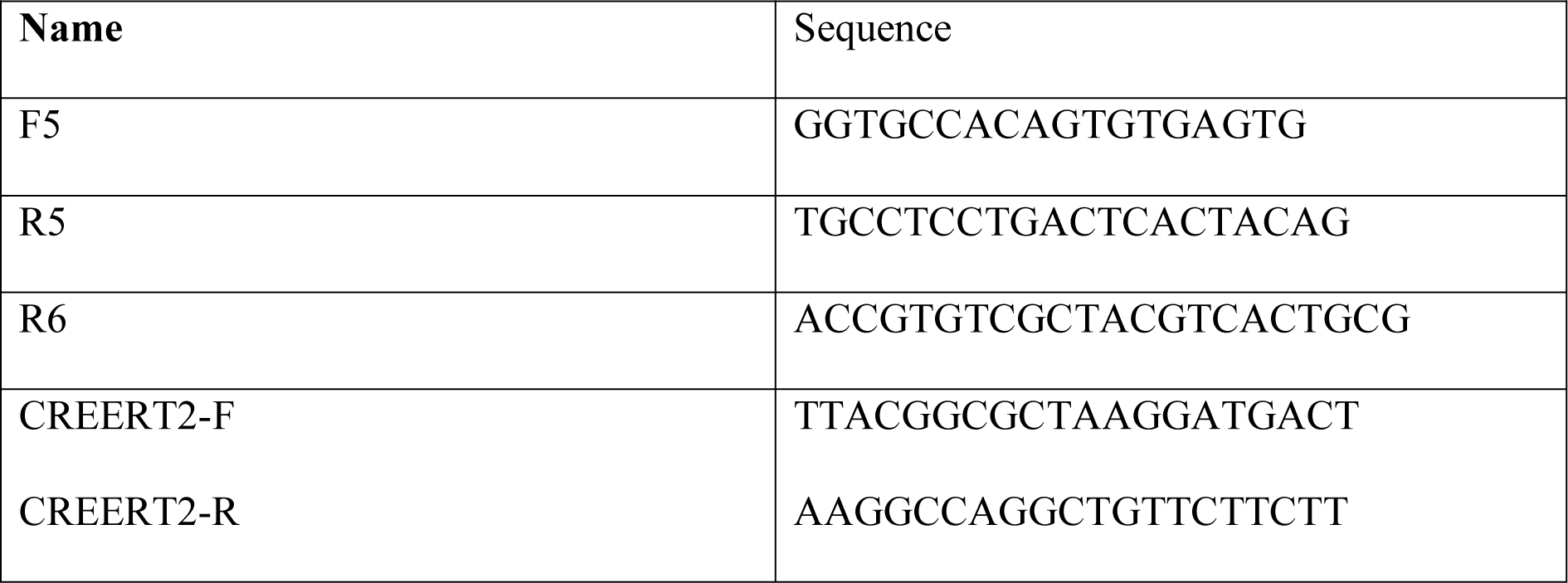

### Statistical analysis

For the outcome endpoints, the disease-free survival (DFS) was defined as the measure of time after treatment during which one of the following events occurred: relapse at local or distant sites, or intercurrent death without recurrence. Overall survival (OS) was defined as the time between surgery and death from any cause. Survival curves were analyzed by the Kaplan-Meier method and compared using the log-rank test. Cox’s proportional hazards model, adjusted for other prognostic factors (i.e., gender, tumor location, tumor grade, tumor stage, and RPAP3 status), was used to evaluate the association of RPAP3 expression with outcome.

Transcriptomic analysis of R2TP subunits in human samples were extracted from the CODREAD cohort available at UCSC Xena platform for public and private cancer genomics data visualization and interpretation https://xenabrowser.net/ (Goldman et al., 2018). One-way ANOVA test was performed using GraphPad Prism 5.0.

Statistical analysis was performed employing the following two statistical software packages: GraphPad Prism 5.0 and SPSS 15.0 (SPSS Inc., Chicago, IL); p < 0.05 was considered as statistically significant. For group comparisons, normality was first tested to choose a pertinent statistical test, as described in the text and Figure legends.

