## supplemental figures and legends, supp. tables for "The R2TP chaperone assembles cellular machineries in intestinal CBC stem cells and progenitors"

#### Supplemental Figure 1

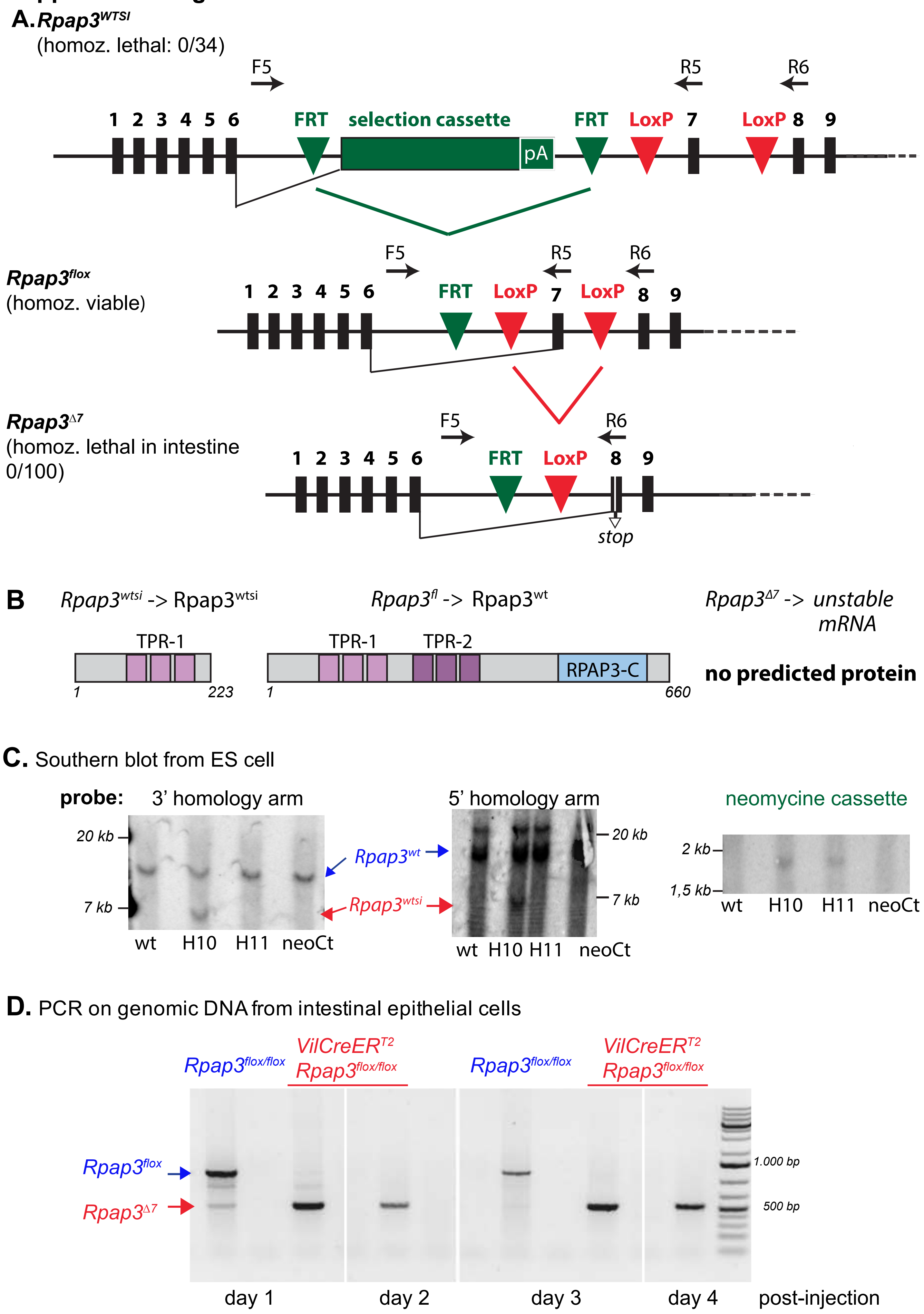

#### Supplemental Figure 2

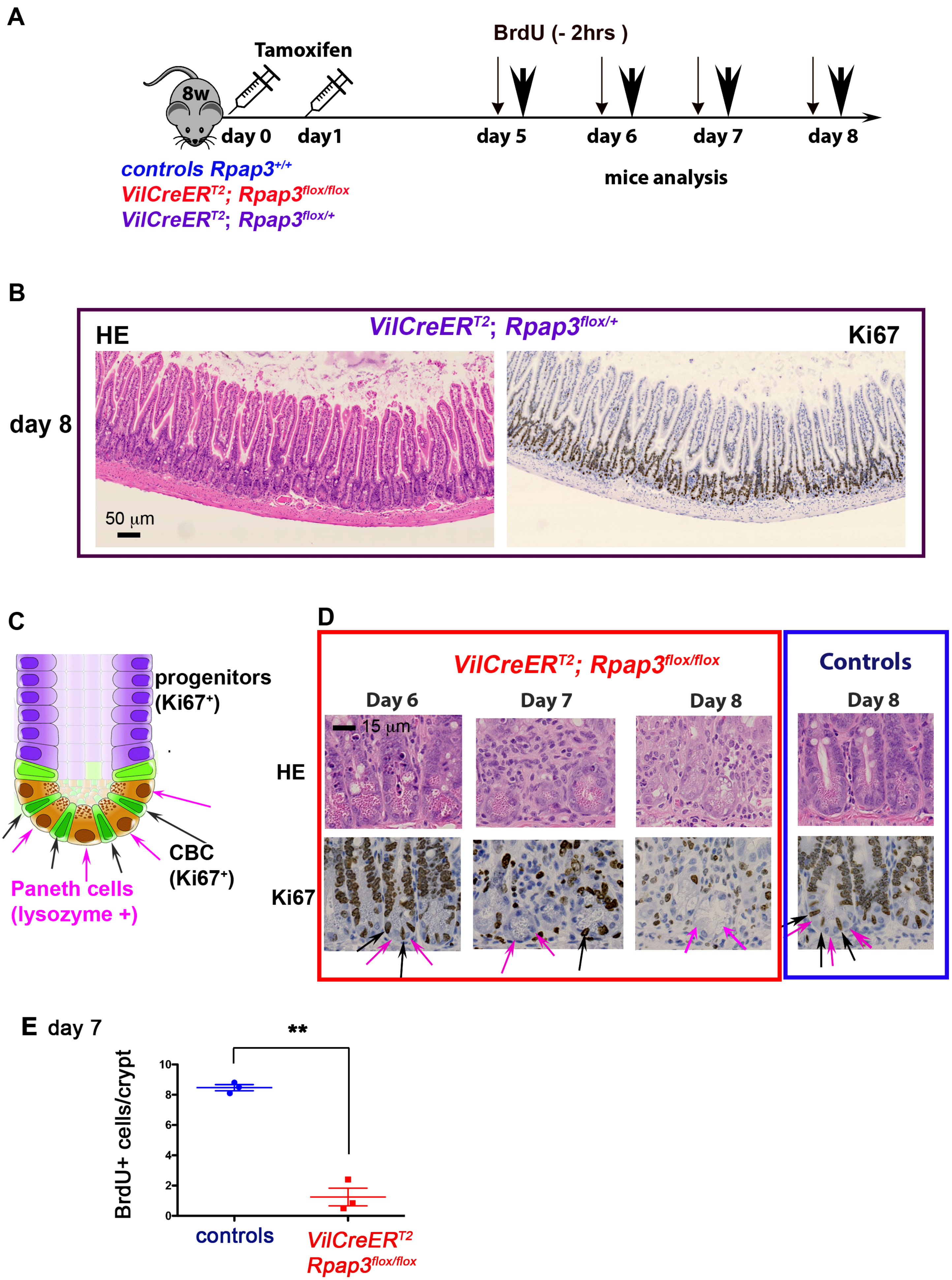

#### Supplemental Figure 3

##### A. Olfm4 at day 7

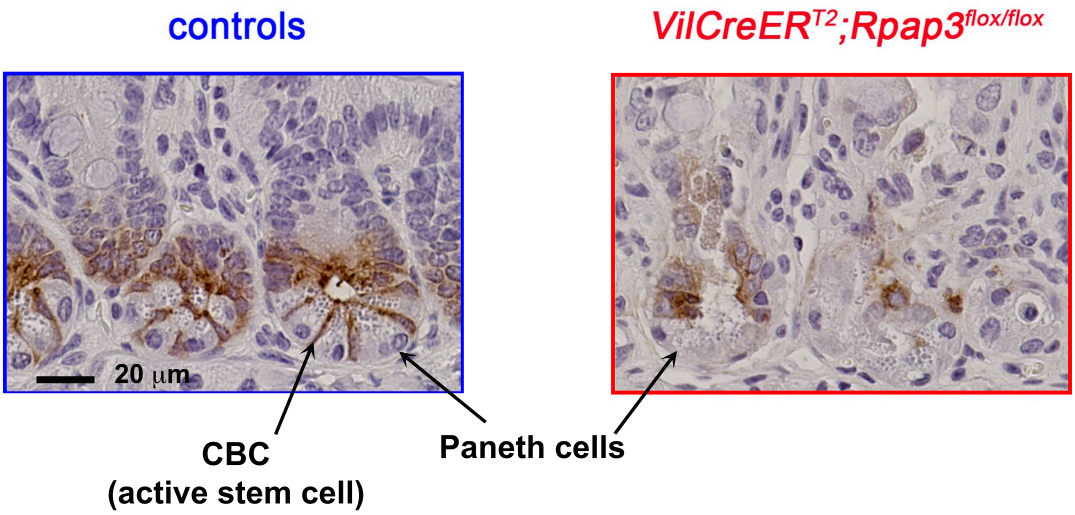


##### B. Western Blot at day 6

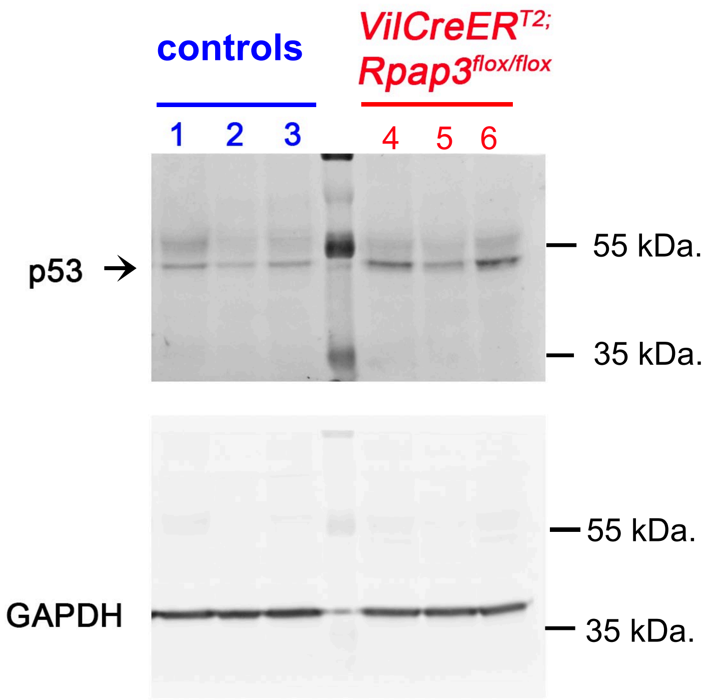

### Supplemental Figure 4

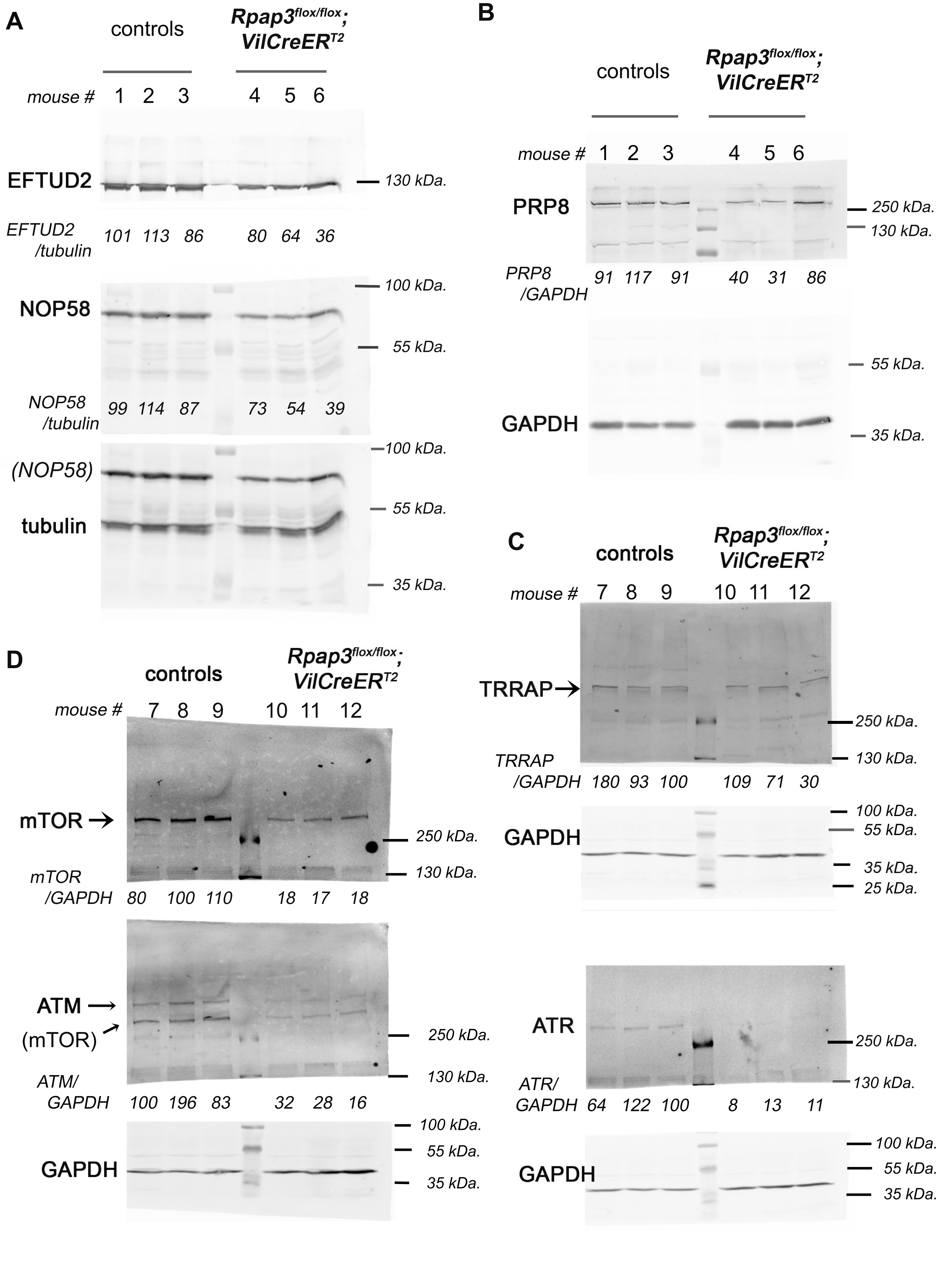

#### Supplemental Figure 5

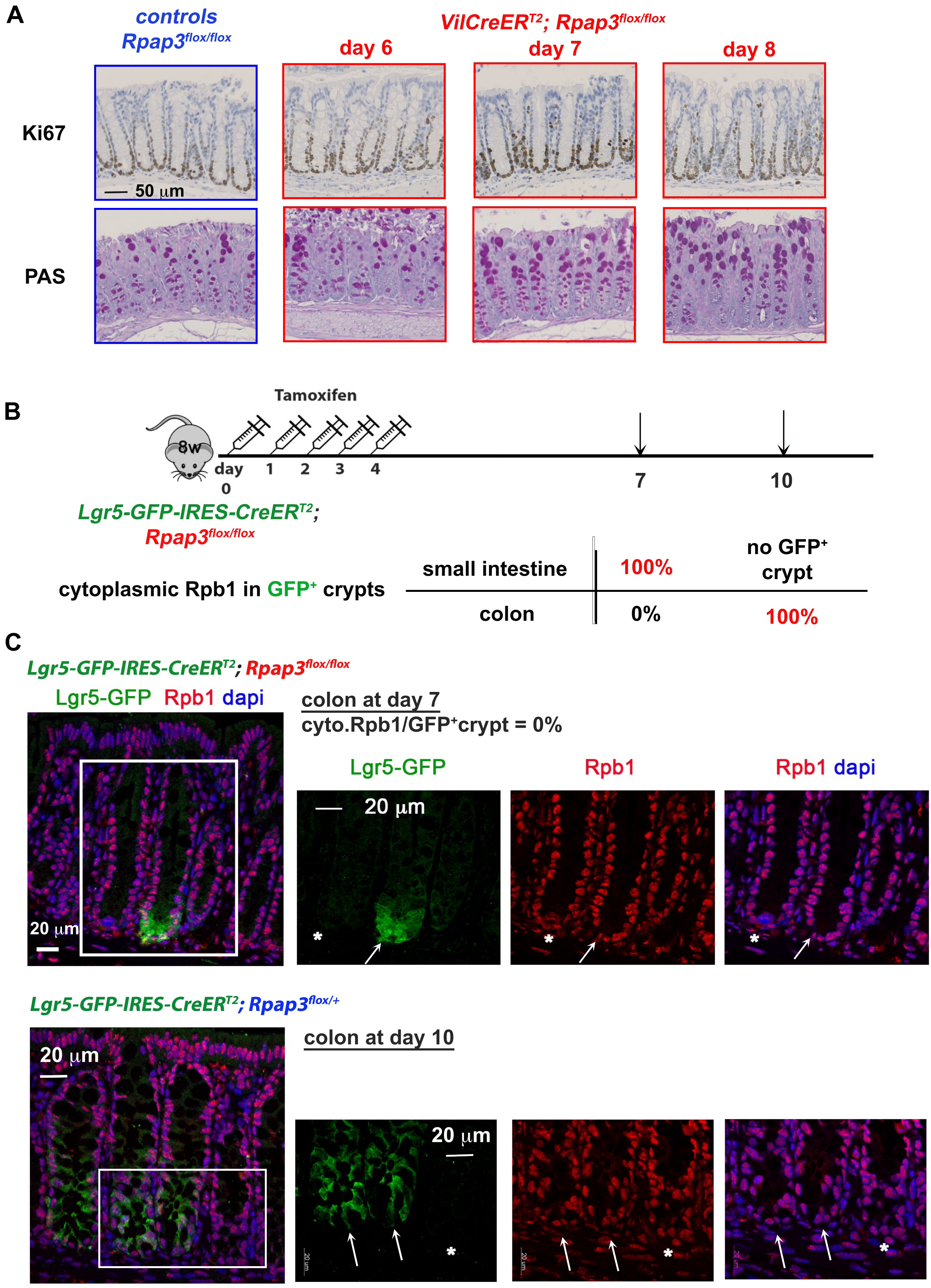

**Supplemental Table 1.** Multivariate analysis of RPAP3expression in CRC tumors (n = 177)

| Variable | HR | 95% CI | P |
|---|---|---|---|
| **Disease-free survival** | | | |
| Gender (Male *vs* Female) | 1.1 | 0.6-2.1 | 0.736 |
| Tumor Location (Rectal *vs* Colon) | 1.9 | 0.9-4.0 | 0.091 |
| Tumor Grade (2-3 *vs* 1) | 1.2 | 0.4-3.8 | 0.823 |
| Tumor Stage (II *vs* I) | 2.3 | 1.2-4.5 | 0.012* |
| RPAP3 (High *vs* Low) | 2.7 | 1.2-6.5 | 0.023* |
| **Overall Survival** | | | |
| Gender (Female *vs* Male) | 1.1 | 0.5-2.2 | 0.885 |
| Tumor Location (Rectal *vs* Colon) | 1.4 | 0.5-3.7 | 0.520 |
| Tumor Grade (2-3 *vs* 1) | 1.2 | 0.3-5.0 | 0.835 |
| Tumor Stage (II *vs* I) | 2.0 | 0.9-4.3 | 0.089 |
| RPAP3 (High *vs* Low) | 2.1 | 0.8-5.5 | 0.133 |

*Statistically significant

**Supplemental Table 2.** Patients and Tumor Characteristics (n = 177)

| **Variable** | **Value (%)** |
|---|---|
| Age at | |
| Median | 70.0 |
| Range | 36 - 90 |
| Gender | |
| Male | 111 |
| Female | 66 |
| Tumor | |
| Colon | 155 (87.6) |
| Rectal | 22 (12.4) |
| Tumor Stage | |
| I | 80 (45.2) |
| II | 97 (54.8) |
| Pathological | |
| Well | 16 ( 9.0) |
| Moderate | 148 (83.7) |
| Poor | 13 ( 7.3) |
| RPAP3 | |
| Low | 54 |
| High | 123 |

**Supplemental Table 3.** RPAP3 status according to clinicopathological features of patients

| Variable | RPAP3 | | |
|---|---|---|---|
| | Low: n (%) | High: n (%) | *P*° |
| Gender | | | |
| Male | 32 (59.3) | 79 (64.2) | 0.613 |
| Female | 22 (40.7) | 44 (35.8) | |
| Tumor Location | | | |
| Colon | 49 (90.7) | 106 (86.2) | 0.4 |
| Rectum | 5 ( 9.3) | 17 (13.8) | 67 |
| Tumor Stage | | | |
| I | 18 (33.3) | 62 (50.4) | 0.0 |
| II | 36 (66.7) | 61 (49.6) | 49* |
| Tumor Grade | | | |
| 1 | 4 ( 7.4) | 12 ( 9.8) | 0.7 |
| 2-3 | 50 (92.6) | 111 (90.2) | 79 |

**Supplemental Table 4: Antibodies used in IHC or IF on paraffin-embedded tissues.**

| **antigen** | **company** | **Clone reference** | **concentration** | **dilution used** |
|---|---|---|---|---|
| cleaved caspase-3 | Cell signaling | 9664 | unknown | 1/2000 |
| Olfm4 | Cell signaling | D6Y5A XP | unknown | 1/500 |
| Rpb1 | Euromedex | 1PB7C2 | unknown | 1/3000 |
| BrdU | Biolegend | 339810 | 0.5mg/ml | 1/100 |
| Ki-67 | Invitrogen | solA15 | 0.5mg/ml | 1/1000 |
| GFP | ThermoFisher | a-6455 | unknown | 1/100 |
| Lysozyme C | Santa Cruz | sc-27958 | unknown | 1/400 |
| RPAP3 | proprietary | 19B11 | unknown | 1/10 |

All antibodies have been validated by the manufacturer. Anti-human RPAP3 has been validated and previously published [16, 39].

### Supplemental Figure legends

**Supplemental Figure 1: Targeting *Rpap3* in the murine intestinal epithelium**

(A) Schematic representation of the different *Rpap3* alleles described in this paper. Black boxes represent the 9 first exons in *Rpap3* murine gene with numbering above. Green and red forms represent the inserted elements : green triangles for FLP recombination sequences, green box represents the selection cassette containing the β-galactosidase ORF and the poly-adenylation site (pA), red triangles indicate the *LoxP* sites. Black arrows represent the primers used for genotyping.

(B) Rpap3 proteins encoded by the above-described alleles with domains and amino acid numbering in italic.

(C) Southern blot of genomic DNA showed a correct and unique recombination of the *wtsi* construct at the *Rpap3* locus in the KOMP ES clone H10 but not in H11. Molecular sizes are indicated on the side of the pictures.

(D) PCR on genomic DNA from small intestine epithelial cells shows that recombination occurs as early as one day after tamoxifen injection in the *VilCreER$_{T2}$; Rpap3$_{flox/flox}$* mice, and persists up to 4 days (amplification of a band corresponding to *Rpap3$_{\Delta 7}$*). PCR on genomic DNA from *Rpap3$_{flox/flox}$* mice intestinal epithelial cells amplifies a band corresponding to the floxed, unrecombined *Rpap3* allele. Molecular sizes are indicated on the right.

**Supplemental Figure 2**: **Invalidation of *Rpap3* compromises the CBCs and progenitors.**

(A) Schematic representation of the experimental setting.

(B) Hematoxylin and Eosin (HE, left panel) and IHC for Ki67 (right panel) on jejunums from *VilCreER$_{T2}$; RPAP3$_{flox/+}$* mice at day 8. Scale bar represent 50µm. Panels are representative from n=4 animals from two independent experiments.

(C) Schematic representation of a crypt from the small intestine. Only dividing CBCs (in green, black arrows) and progenitors forming the Transient Amplifying compartment (TA, purple) are

Ki67+. Paneth cells (identified by a characteristic, punctuated cytoplasm stained by (HE), pink arrows) are Ki67-.

(D) Zoom from HE and Ki67 stainings of jejunums from *VilCreER$_{T2}$; Rpap3$_{flox/flox}$* presented in Figure 3 B. Pink arrows indicate Paneth cells and black arrows CBC stem cells.

(E) Graph shows mean number of BrdU+ cells/crypt in n=4 controls and *VilCreER$_{T2}$; Rpap3$_{flox/flox}$* animals at day 7. Each point represents the average number of BrdU+ cells calculated in n>35 crypts from two different zones per animal. Mean values with S.E.M are indicated for each experimental group. Unpaired two-tailed t test with Welch's corrections (t=11.66, df=2) indicates significant difference between controls and *VilCreER$_{T2}$; Rpap3$_{flox/flox}$* animals (p=0,0073; n=3).

**Supplemental Figure 3**: **Invalidation of *Rpap3* induces p53 stabilization**

(A) Enlarged fields from the images in Figure 3 (A), compare Olfm4 staining in crypts of the jejunum from control and *VilCreER$_{T2}$; Rpap3$_{flox/flox}$* littermates, at day 7. Scale is identical for the two pictures.

(B) Western blot analysis of p53 levels in the lysates of crypt cells from *VilCreER$_{T2}$; Rpap3$_{flox/flox}$* animals, as compared to control littermates. Each lane represents lysates prepared from one animal with the indicated genotype. GAPDH was used as a loading control. Molecular weights are indicated on the right. Animals belong to a different experiment than those used to monitor p53 by IF, presented in Figure 3F.

**Supplemental Figure 4**: **Invalidation of *RPAP3* affects NOP58, PRP8, EFTUD2, PRP8, mTOR, ATR, ATM but not TRRAP expression levels.**

Expanded views of the Western blot presented in Figure 4. (A) Intestinal epithelial crypt cell lysates from 3 controls and 3 *VilCreER$_{T2}$; Rpap3$_{flox/flox}$* mice are shown. Western blot was probed with anti-EFTUD2, anti-NOP58 and tubulin, as a control (note the remains of NOP58 signal upon tubulin detection).

(B) Same as in A, but membranes were probed with antibodies recognizing PRP8 and GAPDH.

(C, D) Same as in A, Western blots were probed against mTOR, TRRAP, ATM (with residual signal for mTOR detection) and ATR. GAPDH was used as a loading control.

(A-D) Quantification of the signal ratios are indicated below each lane (average for the control ratios was arbitrarily set to 100). Molecular weights are indicated on the right.

**Supplemental Figure 5: Cytoplasmic accumulation of Rpb1 in *Rpap3* KO crypts correlates with cellular turnover.**

(A) Immunostaining for Ki67 (top row) and Periodic Aid Schiff staining (PAS, bottom row) of colons from *VilCreER*$^{T2}$*; RPAP3*$^{flox/flox}$ and control littermates from day 6 to 8 after the first tamoxifen injection. Scale bar is identical for all figures. Each panel is representative of n=8-12 animals from three independent experiments.

(B) Experimental setting in the mosaic model *Lgr5-GFP-IRES-CreER*$^{T2}$*; RPAP3*$^{flox/flox}$. In GFP$^{+}$ crypts, Rpb1 accumulates in the cytoplasm of progenitors and CBC stem cells at day 7 in the small intestine and at day 10 in the colon (n=3).

(C) Analysis of the colons from *Lgr5-GFP-IRES-CreER*$^{T2}$*; Rpap3*$^{flox/flox}$ animals at day 7 showed that neither crypts with a detectable GFP signal (white arrows) nor adjacent GFP$^{-}$ crypts (asterisk) displayed Rpb1 accumulation in the cytoplasm (0/19 and 0/21, respectively, in n=3 animals). Colons of *Lgr5-GFP-IRES-CreER*$^{T2}$*; Rpap3*$^{flox/+}$ control animals did not show any detectable cytoplasmic accumulation of Rpb1 (red) in their GFP$^{+}$ crypts (0/17) nor GFP$^{-}$ crypts (0/20).
